# A Chemigenetic Fluorescence Lifetime Biosensor for Dopamine Sensing

**DOI:** 10.64898/2026.09.19.752778

**Authors:** Anh-Thu Nguyen, Hieu Duc Tran, Limin Chen, Abby Criswell, Susanna Bradbury, Yui Zhu, Yujie He, Sohyun Kim, Yuan-I Chen, Trung Duc Nguyen, Soonwoo Hong, Yu-An Kuo, Saeed Seifi, Wei-Ru Chen, Stephanie K. Seidlits, Lief E. Fenno, Hsin-Chih Yeh

## Abstract

Measuring the precise dynamics of specific neuromodulators in neurons is key to understanding how information is transmitted and processed in the brain. Despite the growing use of intensity-based neurochemical sensors such as dLight in recent years, it remains challenging to monitor the dynamics of multiple neurochemicals simultaneously in live neurons. In contrast to intensiometric sensors that are susceptible to artifacts caused by changes in sensor expression and rapid photobleaching, fluorescence lifetime-based sensors are less affected by these factors and can provide more stable measurements. Here we introduce dHaloLife_635_, a chemigenetic fluorescence-lifetime-based dopamine (DA) sensor that exhibits a fluorescence lifetime decrease of 0.21 ns upon dopamine binding. dHaloLife_635_ consists of a G protein-coupled dopamine receptor (hDRD1), a circularly permuted HaloTag (cpHaloTag), and a HaloTag ligand-functionalized cell-permeable dye (JF_635_-HTL). Dopamine binding to dHaloLife_635_ induces a conformational change in the cpHaloTag that alters the local environment of the conjugated dye, thereby changing its fluorescence lifetime. In live HEK293T cells, dHaloLife_635_ responds to dopamine but not to serotonin, L-DOPA, or GABA, and only marginally to norepinephrine, with selectivity for dopamine over norepinephrine confirmed in primary neurons, demonstrating excellent specificity. Together, these results establish dHaloLife_635_ as an initial step toward a broader family of chemigenetic lifetime sensors that exhibit distinct fluorescence lifetime signatures upon binding their respective analytes, thereby potentially enabling highly specific and sensitive monitoring of multiple neurochemical dynamics *in vivo* over extended timescales.

## INTRODUCTION

Neurotransmitters (NTs) and neuromodulators (NMs) are chemical messengers that play essential roles in communication among neurons in the brain. More than 100 NT and NM molecules with either known or putative physiological actions in the brain have been identified, encompassing amino acids, monoamines, nucleotides, neurolipids, and neuropeptides^1,2^. Precisely measuring the dynamics of a specific neurochemical is therefore critical for understanding how information is transmitted and processed in the brain and how these molecules contribute to neurological and psychiatric disorders^3,4^. Neurochemicals often act by binding to G-protein-coupled receptors (GPCRs), initiating molecular signaling cascades that regulate synaptic strength, neuronal excitability, and neural circuit dynamics over timescales ranging from tens of milliseconds to hours^5,6^. To measure these dynamics *in vivo*, genetically encoded GPCR-based intensiometric sensors, whose fluorescence intensity changes upon neurochemical binding, have been developed and widely adopted in neuroscience in recent years because of their high spatial (cellular or subcellular) and temporal (millisecond) resolution, broadly tunable sensing concentration range (pM-mM), and excellent molecular specificity^4^. These GPCR-based sensors are constructed by inserting a circularly permuted fluorescent protein (cpFP) into the third intracellular loop (ICL3) of a GPCR^7^, located between the 5^th^ and 6^th^ transmembrane domains^4^ (TM5 and TM6). Upon neurochemical-binding-induced GPCR activation, ICL3 undergoes a conformational change^8–10^ (**Supplementary Figure S1**) that alters the local chromophore environment of the cpFP, thereby changing its fluorescence intensity^7^. This design strategy has been employed to develop two GPCR-based sensor series – the Light family (e.g., dLight, nLight and psychLight) established by Tian’s group and others^4,7^ and the GRAB family (e.g., GRAB_DA_ and GRAB_ACh_) established by Li’s group^11,12^, enabling *in vivo* sensing of a wide range of neurochemicals, including dopamine (dLight^7^ and GRAB_DA_^11^), acetylcholine (GRAB_ACh_^12,13^), norepinephrine (nLight^14^ and GRAB_NE_^15^), and serotonin (psychLight^16^ and GRAB_5-HT_^17^). While these sensors have enabled robust detection of physiologically and behaviorally relevant neurochemical transients that govern rapid changes in brain state, the sensor signals are influenced not only by changes in neurochemical concentrations but also by factors such as tissue absorption/scattering^18^, photobleaching^19^, fluctuations in excitation power^20^, spectral bleedthrough^21^, variations in sensor expression levels^18,22^ and artifacts arising from autofluorescence, hemodynamics, and animal movement^23^, making it challenging to precisely characterize the absolute concentration of a specific neurochemical in a given brain region over timescales of hours to days.

To address the limitations of intensiometric sensors, researchers have started using GPCR-based sensors whose fluorescence lifetime (an intrinsic photophysical property of fluorophores) changes upon target neurochemical binding, such as GRAB_ACh3.0_ for acetylcholine (ACh) sensing^24^ and dLight3.8 for dopamine (DA) sensing^23^. Compared with intensiometric measurements, fluorescence lifetime measurements are not only insensitive to measurement artifacts, sensor expression variations and photobleaching^25,26^, but also allow absolute analyte concentrations to be stably acquired, thus enabling precise comparisons across imaging days, samples and animals^24,27^. While fast (milliseconds to seconds) and slow (hours to days) neuronal signals have been obtained in live animals using the lifetime-based sensors GRAB_ACh3.0_ and dLight3.8^23,24^, these sensors have only moderate brightness and photostability and are generally limited to the green to red spectral range (500-620 nm) due to the use of common fluorescent proteins as reporters^2^. Whereas researchers have attempted to develop GPCR-based sensors with far-red or near-infrared (NIR) emission (>650 nm) to support multiplexed detection, it has proven very challenging to obtain suitable and sufficiently bright circularly permuted far-red/NIR fluorescent proteins^28,29^. Replacing the FP reporters with bright, photostable, and cell-permeable dyes with far-red emission^27,30^ would both further enhance the *in-vivo* performance of neurochemical sensors through the dyes’ high brightness and photostability^31,32^ and open the door to simultaneous monitoring of multiple neurochemicals in live animals^30^.

To address the limitations of FP reporters, a chemigenetic sensor using a far-red dye as the reporter, termed HaloDA1.0, has been demonstrated by Li’s group for dopamine (DA) sensing^30^. However, despite its large intensity increase upon DA binding, no lifetime changes were reported for HaloDA1.0. Although another lifetime-based chemigenetic sensor with far-red emission, termed WHaloCaMP, has been developed by Farrants and Schreiter for Ca^2+^ imaging^27^ and exhibits a remarkable +2.1 ns lifetime change upon calcium binding, it is not a GPCR-based sensor for neurochemical detection. Inspired by these works, here we demonstrate the design of a new class of chemigenetic sensors that are (1) GPCR-based, (2) far-red-emitting, and (3) capable of showing a measurable fluorescence lifetime change upon target neurochemical binding. Consisting of a dopamine receptor (hDRD1, a GPCR) fused with a circularly permuted self-labeling protein tag (cpHaloTag), and a ligand-functionalized cell-permeable dye (JF_635_-HTL; HTL: HaloTag ligand, which is a reactive chloroalkane linker), our dHaloLife_635_ (<u>d</u>opamine <u>Halo</u>Tag- and <u>Life</u>time-based sensor using JF<u>_635_</u> as the reporter) is the first GPCR-based sensor that employs a far-red dye as the reporter while also showing a fluorescence lifetime change (−0.21 ns) upon DA binding. When tested in HEK293T cells and primary neurons, dHaloLife_635_ exhibits dose-dependent changes in both lifetime and intensity, similar to the cpFP-based DA sensors dLight3.6 and dLight3.8^33^. Interestingly, as both an intensiometric and a lifetime-based sensor, dHaloLife_635_ exhibits different dose responses in intensity and lifetime measurements, with half-maximal effective concentrations (EC_50_) of 190 nM and 880 nM for the intensity and lifetime readouts in HEK293T cells, respectively. Compared with dLight3.8, which shows a comparable DA sensing concentration range (1 nM to 100 μM) and lifetime change upon DA binding (+0.24 ns), dHaloLife_635_ offers a potential photostability advantage through its use of a synthetic fluorophore^34^. Moreover, dHaloLife_635_ showed reliable DA sensing in primary neurons. To the best of our knowledge, dHaloLife_635_ is the first far-red-emitting, lifetime-responsive chemigenetic DA sensor based on a HaloTag and GPCR scaffold, distinguishing it from intensity-based chemigenetic DA sensor HaloDA1.0^30^ and the cpFP-based DA sensor dLight3.8^33^.

## RESULTS

### Development and engineering of the dHaloLife_635_ lifetime dopamine sensor

Current intensiometric sensors are mostly based on circularly permuted green fluorescent protein (cpGFP), whose chromophore ionization state depends on the surrounding environment^35^. Similar to GFP, many cell-permeable rhodamine derivatives are also environment sensitive, residing in an equilibrium between a nonfluorescent lactone (L) form and a fluorescent zwitterionic (Z) form^36,37^; these include a number of far-red Si-containing JF dyes (e.g., JF_635_ and JF_646_). The hydrophobic, nonfluorescent lactone form is more cell-permeable than its zwitterionic counterpart, and binding of an L-Z dye (e.g., JF_585_) to a self-labeling protein (e.g., HaloTag) usually shifts the equilibrium toward the fluorescent zwitterion form, creating fluorogenic probes that light up more than 1,000-fold upon binding cellular protein targets^36^. This L-Z equilibrium shift has been further exploited by Lavis, Schreiter, and Li to create chemigenetic calcium sensors^32^, voltage indicators^32^, and dopamine sensors^30^, in which Ca^2+^ binding, membrane potential change, or dopamine binding pushes the JF dyes further toward the zwitterion form, giving rise to a fluorescence increase. Interestingly, in chemigenetic sensor designs, the analyte receptor or analyte-binding domain does not sit in immediate proximity to the dye, suggesting that the environmental change around the dye results primarily from conformational changes transmitted through the linkers^32^.

Inspired by these prior works, we began building our lifetime-based dopamine sensor by replacing the cpFP in dLight1.1^7^ with a circularly permuted HaloTag (cpHaloTag) protein derived from the chemigenetic calcium sensor HaloCaMP^32^. Specifically, the cpHaloTag scaffold was inserted into the third intracellular loop (ICL3) of the human dopamine D1 receptor (hDRD1) (**Supplementary Figures S1** and **S2**). Position 143 within the loop of the HaloTag was chosen as the site for circular permutation to create new N and C termini, given its close proximity to the bound dye and strong retention of functionality^32^ (**Figure 1A**). The sensor construct is completed when the hDRD1-cpHaloTag fusion receptor is labeled with a dye-HTL conjugate (**Figure 1B**). When DA binds the fusion receptor, a large conformational change around the bound dye should shift its equilibrium between the nonfluorescent lactone (L) form and the fluorescent zwitterion (Z) form (**Figure 1C**), leading to a fluorescence intensity or lifetime change. Our goal here was to identify the variant with the largest lifetime change upon DA binding.

**Figure 1.**
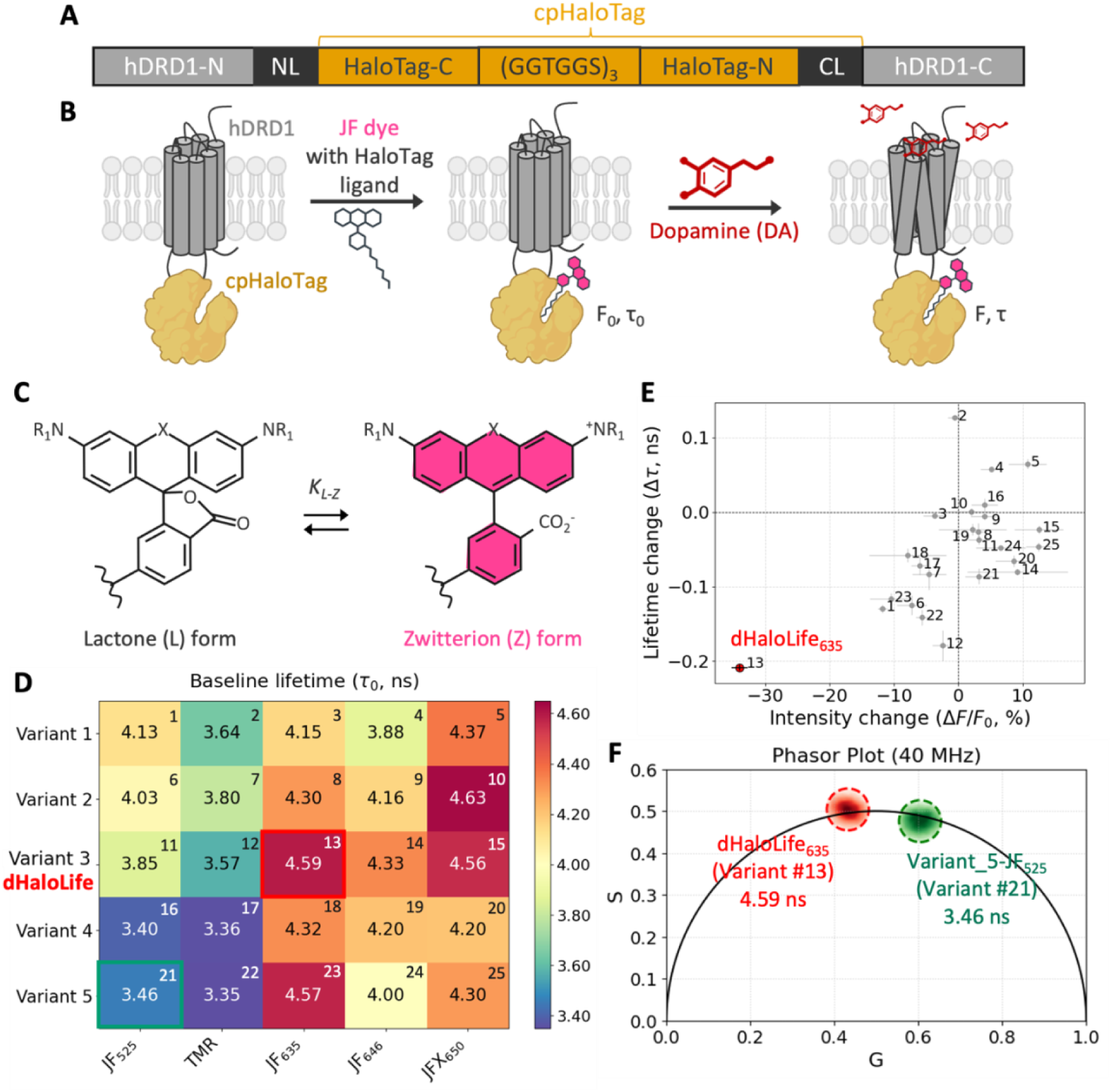
Development of lifetime-based chemigenetic dopamine sensors. (**A)** Design of the dopamine sensors based on human DRD1 (hDRD1) and circularly permuted HaloTag (cpHaloTag, split into HaloTag-C and HaloTag-N subunits). NL and CL denote the N-terminal and C-terminal linkers, respectively. **(B)** Schematic of the chemigenetic dopamine (DA) sensors. A cpHaloTag is inserted into the third intracellular loop (ICL3) of hDRD1 and then conjugated with a Janelia Fluor (JF) dye. Upon DA binding, the sensor undergoes a conformational change that alters the dye’s local environment, thereby changing both its fluorescence intensity and lifetime. Schematics were created with BioRender. **(C)** Fluorogenic mechanism of the rhodamine-based dyes. The dye exists in an environment-sensitive equilibrium (*K_L-Z_*) between the non-fluorescent lactone (L) form and the fluorescent zwitterion (Z) form. **(D)** Baseline fluorescence lifetimes (*τ_0_*) of the 25 sensor variants across 5 hDRD1-cpHaloTag designs and 5 rhodamine derivatives. Variant_3 conjugated with JF_635_ (sensor #13) has one of the longest *τ_0_* (4.586 ± 0.013 ns) and is highlighted in red, while variant_5-JF_525_ (sensor #21) has one of the shortest *τ_0_* (3.459 ± 0.070 ns) and is highlighted in green. **(E)** Scatter plot showing the responses of all 25 sensors upon DA binding. Variant_3-JF_635_ (sensor #13) exhibits the largest fluorescence lifetime response (*Δτ* = *τ* - *τ_0_* = -0.212 ± 0.004 ns) as well as the largest intensity change (*∆F*/*F_0_* = -34.3% ± 1.2%) and is subsequently renamed dHaloLife_635_. **(F)** Phasor plot generated with a digital frequency-domain acquisition module running at a 40 MHz laser repetition rate, showing that the photophysics of the sensors’ reporters (sensor #13 in red and sensor #21 in green) mostly follow a single-exponential decay in the sensors’ unbound states.

Five HaloTag ligand-functionalized rhodamine dyes (JF_525_, TMR, JF_635_, JF_646_ and JFX_650_, **Supplementary Figure S3**) with a range of L-Z equilibrium constants^32,38^ (*K_L-Z_*, **Supplementary Table 1**) were covalently bound to five distinct hDRD1-cpHaloTag fusion receptors (variant_1-5, **Supplementary Table 2**), generating a total of 25 sensor variants (**Figure 1D**). A confocal scanning optical system equipped with a diode laser pulsed at 40 MHz and a digital frequency-domain module^39^ was used for data acquisition and phasor generation (see Methods). When all 25 sensor variants were tested in human embryonic kidney HEK293T cells, hDRD1-cpHaloTag variant_3 using JF_635_ as the reporter (sensor #13, red box in **Figure 1D**) exhibited one of the longest fluorescence lifetimes (4.586 ± 0.013 ns) in its unbound state (the “baseline”, *F_0_* or *τ_0_*). In contrast, variant_5 using JF_525_ as the reporter (sensor #21, green box in **Figure 1D**) showed one of the shortest baselines *τ_0_* (3.459 ± 0.070 ns). Lavis and Schreiter have previously discussed the advantages of high and low *F_0_* in intensiometric sensors – sensors with high *F_0_*are useful for imaging small or sparse sample features, while sensors with low *F_0_* are desirable for imaging in densely labeled samples^32^. Similarly, a long *τ_0_*, which also often means high *F_0_* due to limited nonradiative decay pathways, is desirable for imaging densely labeled samples.

Upon adding 100 μM of DA to the HEK293T cultures, hDRD1-cpHaloTag variant_3 using JF_635_ as the reporter (sensor #13 in **Figure 1E**) exhibited the largest fluorescence intensity and lifetime changes upon DA binding (−34.3% ± 1.2% and -0.212 ± 0.004 ns, **Figure 1E**). Although sensors #2, #4 and #5 showed lifetime lengthening upon DA binding, the absolute extent of change was less than that of sensor #13. Sensor #13 was subsequently renamed dHaloLife_635_, for <u>d</u>opamine <u>Halo</u>Tag- and <u>Life</u>time-based sensor using JF<u>_635_</u> as the reporter. While dHaloLife_635_ acts in the opposite direction (lifetime decreases rather than increases upon DA binding), it is a far-red-emitting chemigenetic sensor with high *F_0_*(at least 70% brighter than their cpFP counterpart^32^) and stable lifetime readings in both its bound and unbound forms, suitable for monitoring fast and slow DA dynamics in neurons. The phasors of dHaloLife_635_ (red circle) and variant_5-JF_525_ (sensor #21, green circle) fell on the universal circle, indicating that these JF reporters’ photophysics follow a simple single-exponential decay in the sensors’ unbound states (**Figure 1F**, also see **Supplementary Figures S4** and **S5**).

**Figure 2** shows the design strategies behind the 5 sensor variants, which focus primarily on variations in the NL and CL linkers. As mentioned above, in chemigenetic sensor designs, the environmental change around the dye results primarily from conformational changes transmitted through the linkers^32^ (**Figure 2A**). The cell-permeable dyes selected for testing were the green dye JF_525_, the orange-red dye tetramethylrhodamine (TMR), and the far-red dyes JF_635_, JF_646_ and JFX_650_, which exhibit low to intermediate *K_L_-_Z_* values^27,32,40^ when functionalized with a HaloTag ligand (HTL). JF_525_ and JF_635_ have been employed as reporters in chemigenetic voltage^32,34^ and calcium^32^ sensors, while JF_646_ and JFX_650_ have more recently been used in sensors for animal studies due to their better bioavailability in the central nervous system^30^ (i.e., good blood-brain barrier permeability).

**Figure 2.**
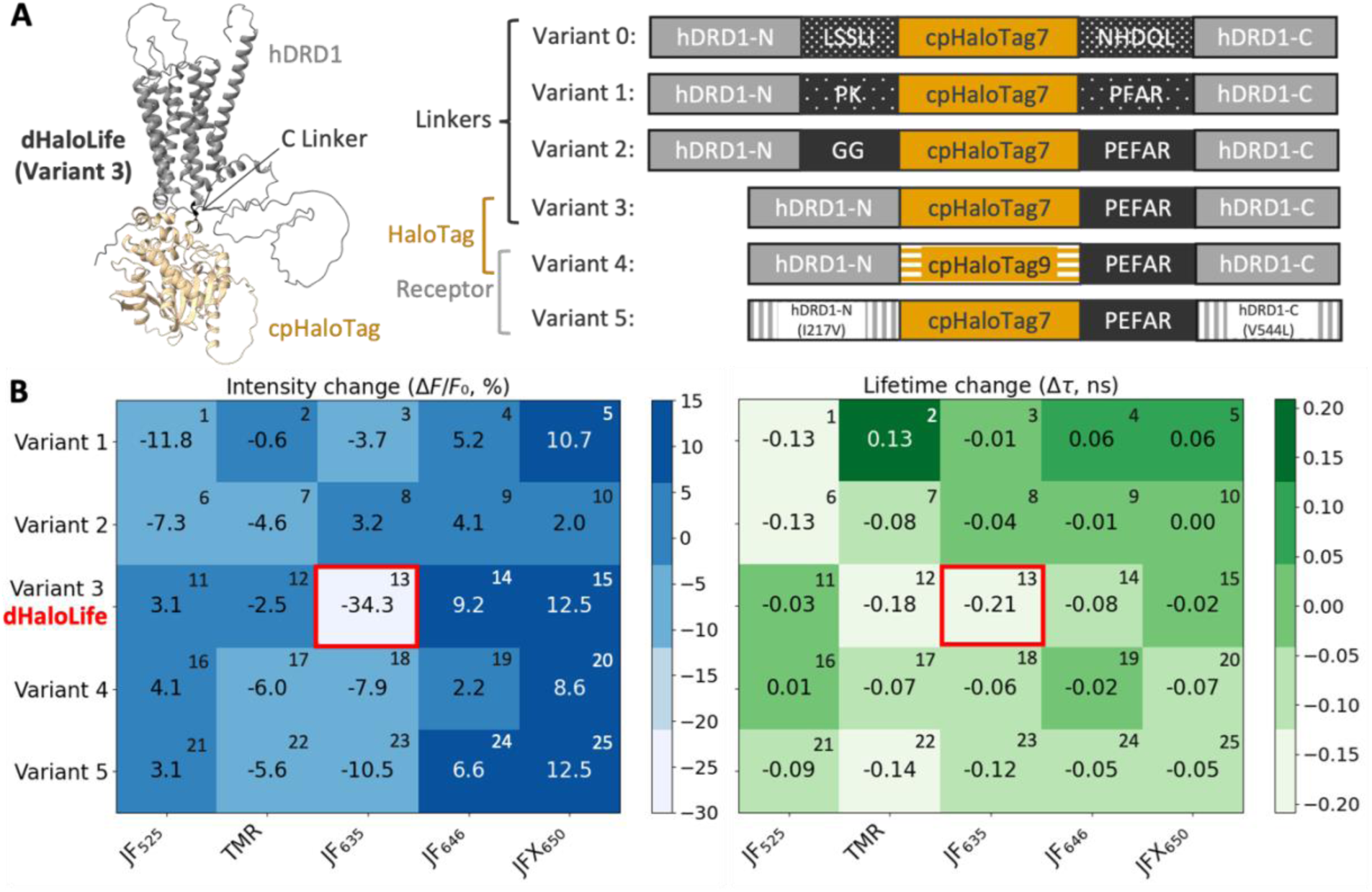
Rational designs of the chemigenetic dopamine sensors. **(A)** (Left) AlphaFold2-predicted 3D structure of dHaloLife (see also **Supplementary Figure S6**). (Right) Structural configurations of linkers and scaffolds for variants_1-variant_5. **(B)** Heatmaps summarizing (Left) the fluorescence intensity changes and (Right) the fluorescence lifetime changes across all 25 sensors in response to 100 µM DA. Intensity response (*ΔF*/*F_0_*, %) was calculated from the mean fluorescence intensity before (*F_0_*) and after (*F*) analyte addition, while fluorescence lifetime response (*Δτ*, ns) was calculated from the average lifetime before (*τ_0_*) and after (*τ*) analyte addition. The red box highlights the top performer (sensor #13, renamed dHaloLife_635_), which exhibits the largest dynamic range in both intensity (−34.3% ± 1.2%) and lifetime (−0.212 ± 0.004 ns). Data represent means with n ≥ 12 cells across ≥ 3 independent experiments.

HEK293T cells were used as the model system to test the 25 sensor constructs. The fluorescence intensity and lifetime were measured before and after the sensor-expressing HEK293T cells were bathed in a saturating concentration of 100 μM DA. Although only 5 engineered sensor variants are shown in **Figure 2**, 17 additional sensor constructs were designed and tested in HEK293T cells; the results are summarized in **Supplementary Tables 2**, **3**, and **4**. In short, most variants exhibited only negligible sensor responses in HEK293T cells. One of our early attempts, termed variant_0, was constructed by simply replacing the cpGFP in dLight1.1^41^ with the cpHaloTag from HaloCaMP1a^32^ (which derives from HaloTag7^42^). Variant_0 therefore shared the same NL (LSSLI) and CL (NHDQL) sequences as dLight1.1^41^. However, this design strategy was unsuccessful; variant_0 showed poor membrane localization after transfection into HEK293T cells.

Variant_1 (**Figure 2B**) was inspired by the HaloCaMP1b and HaloCaMP1a designs^32^, which show excellent calcium responses after conjugation with JF_635_. In variant_1, the NL (PK) and CL (PFAR) sequences were identical to those of HaloCaMP1b^32^. Interestingly, our results indicated that variant_1-JF_525_ (sensor #1) exhibits negative lifetime and intensity responses (*ΔF*/*F_0_* = -11.8 ± 0.7% (mean ± s.e.m.) and *Δτ* = -0.129 ± 0.005 ns), while variant_1-TMR (sensor #2) gives a positive lifetime response (*Δτ* = +0.126 ± 0.003 ns) but no intensity change (*ΔF*/*F_0_* = -0.6 ± 1.1%) upon DA binding.

Different linker designs were explored in variant_2 and variant_3. The variant_2 NL (GG) and CL (PEFAR) sequences were inspired by the GRAB_ACh3.0_ and HaloCaMP1b designs, whereas the variant_3 NL (none) and CL (PEFAR) sequences were designed to test the effect of the NL. Among the 5 dyes, only variant_2-JF_525_ (sensor #6) gave a noticeable response to DA (*ΔF*/*F_0_* = -7.3 ± 2.3% and *Δτ* = -0.125 ± 0.013 ns, **Figure 2B** and **Supplementary Table 3**). By contrast, variant_3-JF_635_ (sensor #13, later renamed dHaloLife_635_) showed the largest response to DA (*ΔF*/*F_0_* = -34.3 ± 1.2% and *Δτ* = -0.212 ± 0.004 ns) as well as good membrane localization. Interestingly, variant_3-TMR also displayed a substantial lifetime response to DA (*Δτ* = -0.179 ± 0.021 ns) but showed almost no intensity change (*ΔF*/*F_0_* = -2.5 ± 1.6%).

Beyond linker variations, HaloTag variants engineered by Johnsson’s group^42^ were also tested. In variant_4, cpHaloTag7 was replaced with cpHaloTag9, the brightest variant in the engineered HaloTag family (HaloTag7-Q165H-P174R), which shows lifetime modulation with TMR and other cell-permeable rhodamine derivatives^42^. Although variant_4 exhibited excellent membrane localization in HEK293T cells, none of its 5 associated sensors gave a substantial response to DA (**Figure 2B** and **Supplementary Table 3**).

Lastly, mutations in the DRD1 receptor were explored. Following the rational design of the dLight series, mutations from dLight1.1 to dLight1.3 (F129A and an inserted 224Q) in the conserved binding pocket were tested in the chemigenetic sensors here. However, the resulting variant yielded suboptimal membrane trafficking. Guided by the dLight1.3b-to-dLight3.8 design (I217V and V544L), we then constructed variant_5, which carried a single mutation (I217V) in hDRD1-N and a single mutation (V544L) in hDRD1-C and maintained robust plasma membrane expression. Although variant_5-TMR (sensor #22) and variant_5-JF_635_ (sensor #23) showed noticeable DA responses (*Δτ* = -0.141 ± 0.010 ns and -0.117 ± 0.008 ns, respectively, **Figure 2B** and **Supplementary Table 3**), they did not outperform dHaloLife_635_ (sensor #13).

### Sensor imaging and characterization in HEK293T cells

To confirm that the observed lifetime change in dHaloLife_635_ was due to DA binding to hDRD1, the selective DRD1 antagonist SCH-23390^41^ (hereafter SCH, 10 µM) was added in the presence of 100 µM DA. As expected, both the intensity and lifetime returned to baseline *F_0_*and *τ_0_* levels (**Figure 3A** and **B**, **Supplementary Figures S7**, **S8**, **Supplementary Tables 5**, **6** for full statistics, and **Supplementary Video 1**). These results demonstrated that the conformational change in the DA receptor, which drives the intensity and lifetime changes of the conjugated dye, is strictly dependent on analyte binding.

**Figure 3.**
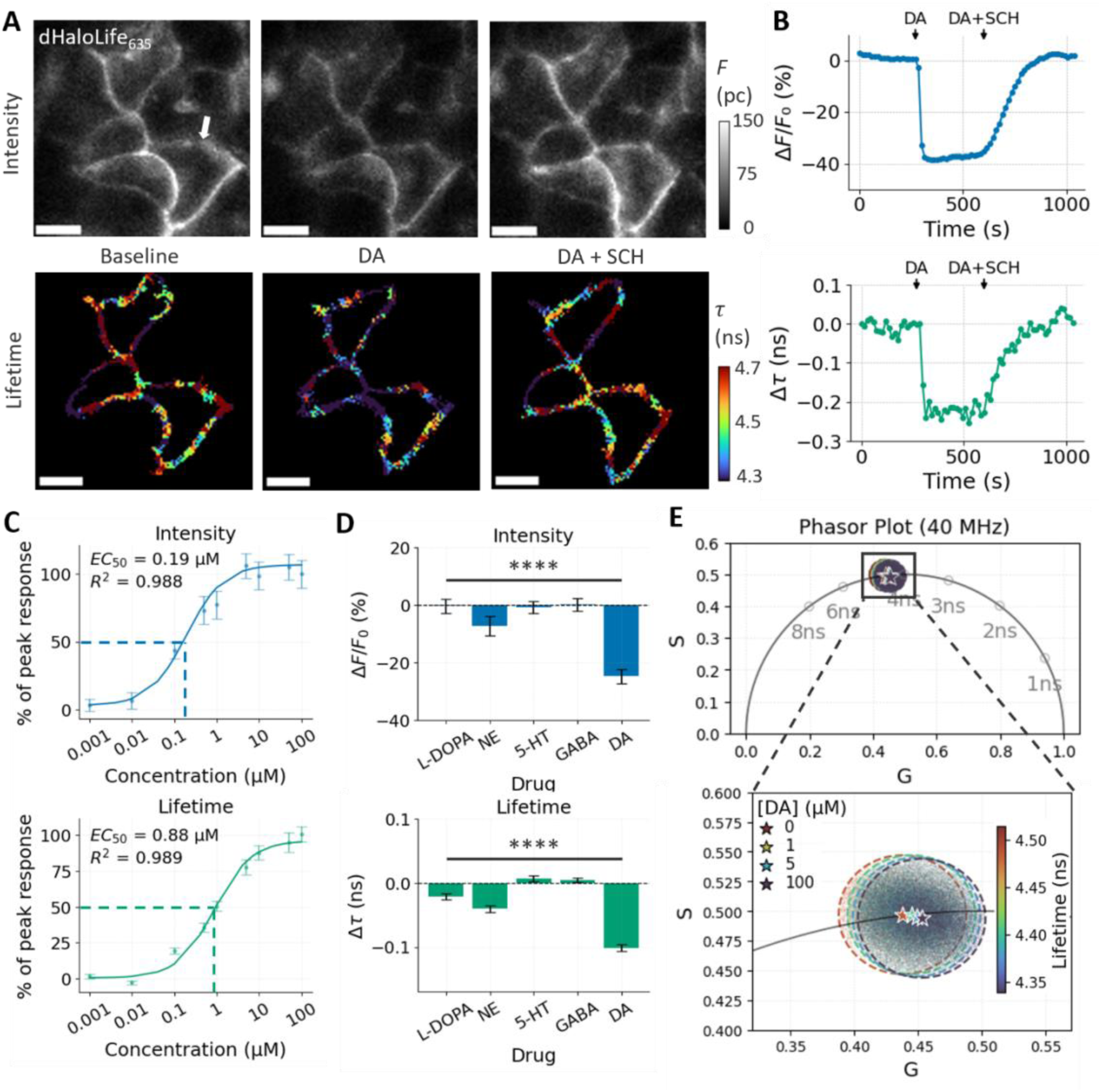
Pharmacological characterization and sensitivity of dHaloLife_635_ in HEK293T cells. **(A)** Representative intensity (top row) and fluorescence lifetime (bottom row) images of HEK293T cells expressing dHaloLife_635_. Cells were imaged at baseline, after addition of 100 µM dopamine (DA), and after 100 µM DA co-application with the DRD1-specific antagonist SCH-23390 (SCH, 10 µM). Scale bars: 10 µm. Intensity was reported as photon counts (pc). **(B)** Representative time traces showing the antagonist-induced reversal of the sensor response in both fluorescence intensity (*ΔF/F_0_*, %, top) and lifetime (*Δτ*, ns, bottom) upon sequential addition of DA and DA+SCH. Time traces represent the response of the denoted cell in (A). **(C)** Dose-response titration of dHaloLife_635_. The sensor exhibits sigmoidal responses to DA with EC_50_ of 190 nM and 880 nM for the intensity and lifetime readouts, respectively. **(D)** Selectivity of dHaloLife_635_ against structurally related neurochemicals at 1 µM. Significant responses are observed only for DA, with marginal cross-reactivity to norepinephrine (NE). **** p < 0.0001, using Brown-Forsythe and Welch ANOVA tests. Data represent the mean ± s.e.m. from n ≥ 14 cells across ≥ 3 independent experiments. **(E)** Phasor plot trajectory showing the transition of the lifetime population across increasing DA concentrations (0, 1, 5, and 100 µM).

To evaluate whether dHaloLife_635_ can be used for quantitative DA imaging, we tested the sensor against different concentrations of DA (1 nM to 100 µM). Consistent with the intensity-based measurements, the lifetime of dHaloLife_635_ showed a robust, dose-dependent decrease (**Figure 3C**). Interestingly, the lifetime readout was less sensitive to DA than the intensity readout, with half-maximal effective concentrations (EC_50_) of 190 nM and 880 nM for the intensity and lifetime readouts in HEK293T cells, respectively. Chen’s group reported an analogous divergence between the two readouts in their cpFP-based GRAB_ACh3.0_ sensor^24^ (EC_50_ of 1,300 nM and 240 nM for the intensity and lifetime readouts, respectively, for ACh sensing^24^), although in that case the lifetime readout was the more sensitive of the two. This likely reflects the distinct underlying mechanisms governing fluorescence intensity and lifetime.

Specificity is critical for applying dHaloLife_635_ in complex neurobiological systems, so we next examined its molecular specificity. dHaloLife_635_ was tested against a panel of related neurochemicals and precursors, including levodopa (L-DOPA), γ-aminobutyric acid (GABA), norepinephrine (NE), serotonin (5-HT), and dopamine (DA), each at 1 µM (**Figure 3D**). The sensor showed no significant response to L-DOPA, GABA, or 5-HT, confirming its high specificity. Because NE is structurally similar to DA, dHaloLife_635_ showed a marginal response, but this was significantly smaller than the response to DA at the same concentration. These data indicated that dHaloLife_635_can selectively report DA in the presence of diverse neurochemicals (****p < 0.0001 for both intensity and lifetime readouts, using Brown-Forsythe and Welch ANOVA tests, **Figure 3D**, see **Supplementary Tables 7** and **8** for full statistics).

To further validate the sensor response, we engineered a control construct by introducing a D103A mutation in dHaloLife to abolish DA binding, termed dHaloLife-mut, following the same strategy used for dLight1.1 validation^7^ (**Figure 4A**). As expected, both the intensity and the lifetime of dHaloLife-mut_635_ showed negligible responses to 100 µM DA (**Figure 4B-C, Supplementary Figure S9**, and **Supplementary Video 2**). The mutant construct conjugated with JF_525_, TMR, JF_635_, JF_646_ and JFX_650_ also showed no response to DA (**Figure 4D** and **Supplementary Tables 3** and **4**). These results confirmed that DA binding is essential for the photophysical changes in dHaloLife sensors.

**Figure 4.**
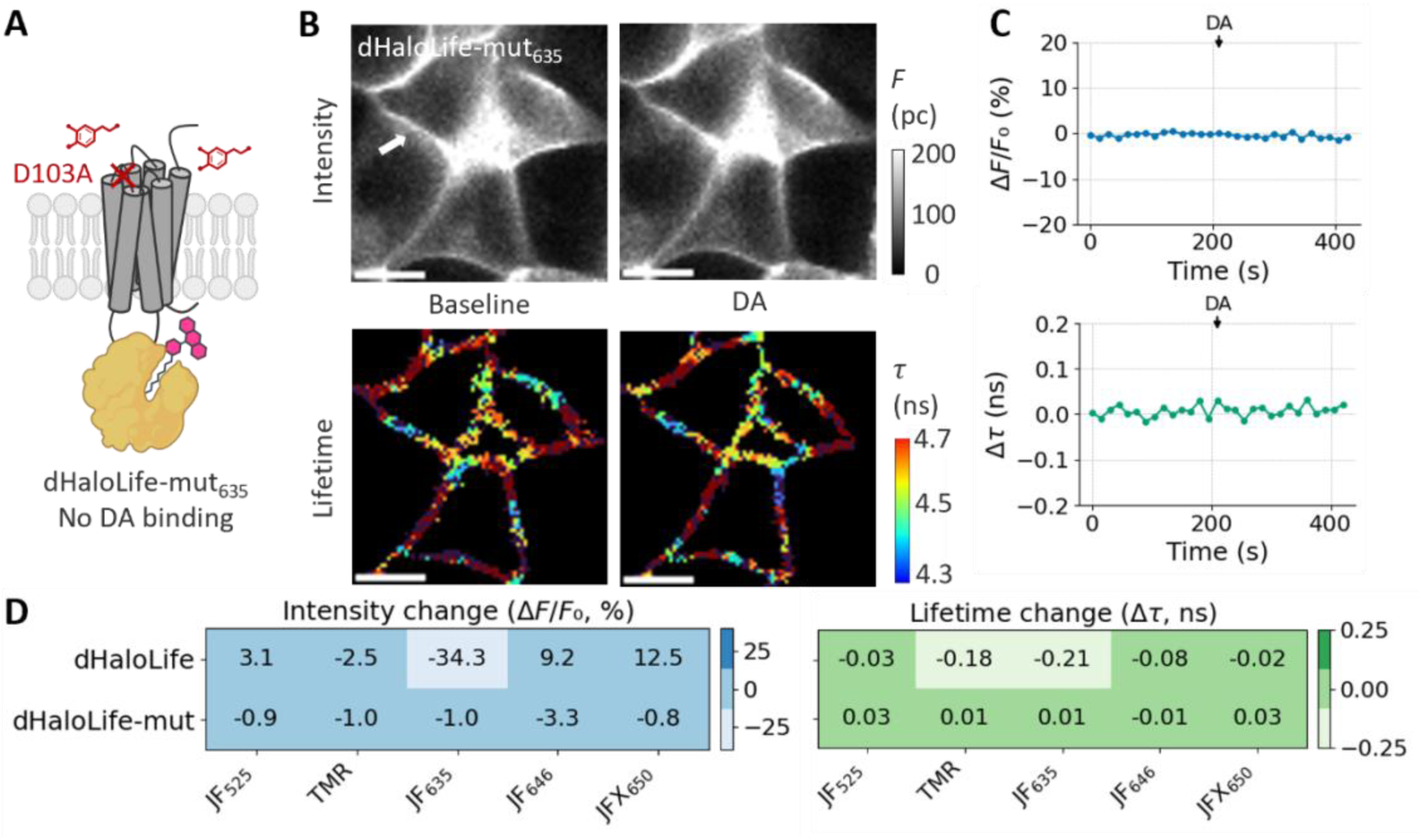
Validation of the non-responsive control sensor, dHaloLife-mut. **(A)** Schematic of dHaloLife-mut_635_ featuring the D103A mutation to abolish dopamine binding. **(B)** Representative intensity and lifetime images of HEK293T cells expressing dHaloLife-mut_635_ showing no response to 100 µM DA. Scale bars: 10 µm. **(C)** Time traces of intensity (*ΔF/F_0_*, %, top) and lifetime (*Δτ*, ns, bottom) confirming the absence of ligand-induced modulation. Time traces represent the response of the denoted cell in (B). **(D)** Heatmaps showing negligible intensity and lifetime changes across all tested rhodamine dyes, validating the structural integrity of the mutant scaffold. Data represent the mean ± s.e.m. from n ≥ 15 cells across ≥ 3 independent experiments.

### Advantages of lifetime over intensity for dopamine sensing Robustness against heterogeneous expression levels

Fluorescence intensity readouts are inherently influenced by sensor expression level, which often varies substantially across cells or tissue regions. Consistent with this, baseline fluorescence intensity of dHaloLife_635_ in HEK293T cells was broadly distributed (median = 29 photon counts (pc), interquartile range (IQR) from 18 to 97 pc, coefficient of variation (CV) = 92.27%, n = 81 cells from 16 independent experiments), and 100 µM DA reduced it to a median of 22 pc (IQR from 11 to 69 pc, CV = 93.48%, ****p < 0.0001, paired Student’s t-test, **Figure 5A**, left). In contrast, the lifetime of dHaloLife_635_ was tightly distributed at baseline (median = 4.530 ns, IQR from 4.490 to 4.678 ns, CV = 2.57%) and decreased to 4.348 ns (IQR from 4.264 to 4.493 ns, CV = 2.75%) with 100 µM DA (****p < 0.0001, paired Student’s t-test, **Figure 5A**, right).

**Figure 5.**
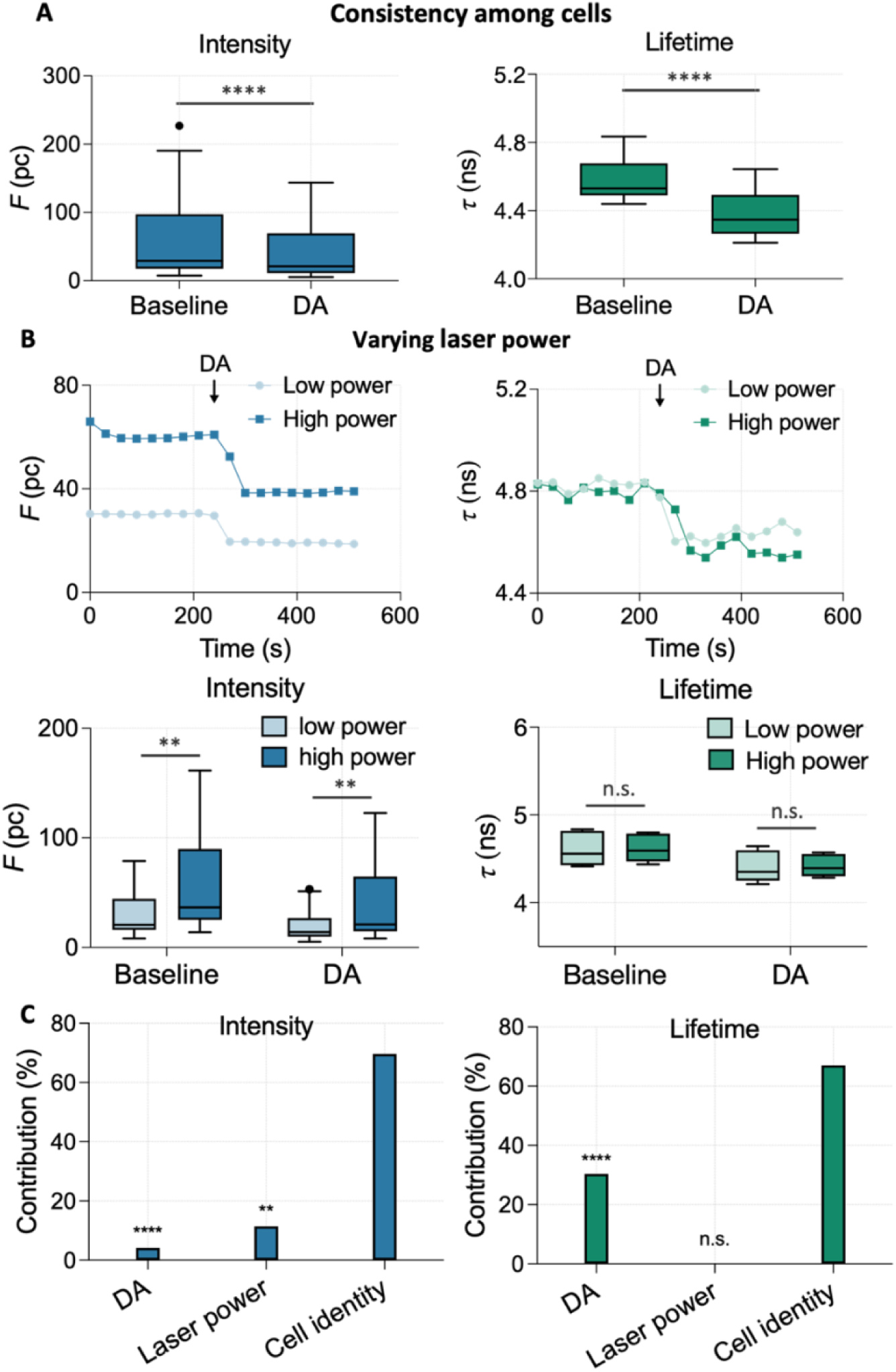
Advantages of fluorescence lifetime readouts over intensity readouts. **(A)** Comparison of baseline and DA-stimulated intensity and lifetime readouts across multiple cells. Whereas intensity measurements could be confounded by heterogeneous expression levels, the absolute lifetime readouts varied less among cells. ****p < 0.0001, two-tailed paired Student’s t-test. Data represent the median ± interquartile range (IQR) with whiskers showing 1.5*IQR from n = 81 cells across 16 independent experiments. **(B)** Invariance to excitation power. Whereas absolute photon counts (*F*) differed markedly between low and high power, the fluorescence lifetime (*τ*) showed no detectable changes across excitation powers. **p < 0.01, and n.s. (not significant) for p ≥ 0.05, two-way ANOVA. Data represent the median ± IQR with whiskers showing 1.5*IQR from n = 18 cells across 4 independent experiments. **(C)** Contribution to total variance of varying laser power test. Two-way ANOVA analysis from (B) showing the contribution of DA stimulation, laser power and cell identity to total variance. ****p < 0.0001, **p < 0.01 and n.s. (not significant) for p ≥ 0.05.

### Fluorescence lifetime of dHaloLife_635_ is unaffected by varying laser power

One of the major challenges in quantitative intensity-based imaging is the dependence of fluorescence signals on excitation power. To examine this effect, we measured the fluorescence intensity and lifetime of dHaloLife_635_ in HEK293T cells across two excitation laser powers. As demonstrated in **Figure 5B**, changes in laser power (from low power to high power, which is approximately 2.5 times stronger in laser power) resulted in a significant shift in the fluorescence intensity (*F*) time traces for the same cells (estimated 220% increase for the shown cell, **Figure 5B**, top). Across cells, higher laser power significantly increased intensity at baseline (median from 21 (low power) to 37 (high power) pc, 95% confidence interval (CI) from 20 – 42 pc (low power) to 34 – 84 pc (high power), **p < 0.01, two-way ANOVA test) and with 100 µM DA (median from 14 to 21 pc, 95% CI from 13 – 27 pc to 22 – 61 pc, **p < 0.01, **Figure 5B**, bottom left). In contrast, lifetime did not differ between laser powers at baseline (median are 4.557 ns (low power) and 4.594 ns (high power), 95% CI from 4.507 – 4.680 ns (low power) and 4.536 – 4.678 ns (high power), non-significant (n.s.)) or with DA (median are 4.352 ns (low power) and 4.394 ns (high power), 95% CI from 4.313 – 4.481 ns (low power) and 4.354 – 4.472 ns (high power), non-significant (n.s.), **Figure 5B**, bottom right). Thus, although absolute intensity depended on laser power, the DA-induced changes calculated for each cell did not: neither the relative intensity change (Δ*F/F_0_*, median -36.1% at low power vs. -36.4% at high power) nor the lifetime shift (Δ*τ* = −0.205 ns vs. −0.200 ns) differed between powers (n.s., two-tailed paired Student’s t-test, n = 18 cells from 4 independent experiments, **Supplementary Figure S10**).

To compare the sources of variation directly, we partitioned the variance of each readout among DA stimulation, laser power, and cell identity (**Figure 5C**). DA stimulation explained approximately 4.1% of intensity variance but 30.3% of lifetime variance (****p < 0.0001), so the sensor response is a much larger share of the lifetime signal than of the intensity signal. Cell identity was the largest contributor for both readouts (69.6% intensity, 67.0% lifetime), reflecting the cell-to-cell differences that remain in either measurement. Most importantly, laser power accounted for approximately 11.5% of intensity variance (**p < 0.01) but no significant fraction of lifetime variance (0.17%, n.s.). These results demonstrate that the absolute lifetime values and the DA-induced lifetime shifts (*Δτ*) of dHaloLife_635_ are consistent across varying laser power settings, ensuring reliable and reproducible readouts across different imaging sessions and microscopy platforms.

### Sensor characterization in primary neuron cells

To evaluate the functional performance of dHaloLife_635_ in a neurobiological context, we expressed dHaloLife in cultured primary hippocampal neurons and characterized sensor intensity and lifetime responses to DA (**Figure 6**). dHaloLife_635_ maintained excellent plasma membrane localization in neurons, as in HEK293T cells (**Figure 6A**). Upon bath application of 100 µM DA, the sensor showed a robust decrease in both fluorescence intensity and lifetime (*ΔF/F_0_* = -30.5 ± 2.8% (mean ± s.e.m.); *Δτ* = -0.167 ± 0.007 ns; **Figure 6A-B**, **Supplementary Figure S11**, and **Supplementary Video 3**). To confirm that these responses were mediated specifically by the DA-binding site of hDRD1, we co-applied the selective antagonist SCH (10 µM), which completely abolished the DA-induced changes, restoring both fluorescence intensity and lifetime to baseline levels (**Figure 6A-B**).

**Figure 6.**
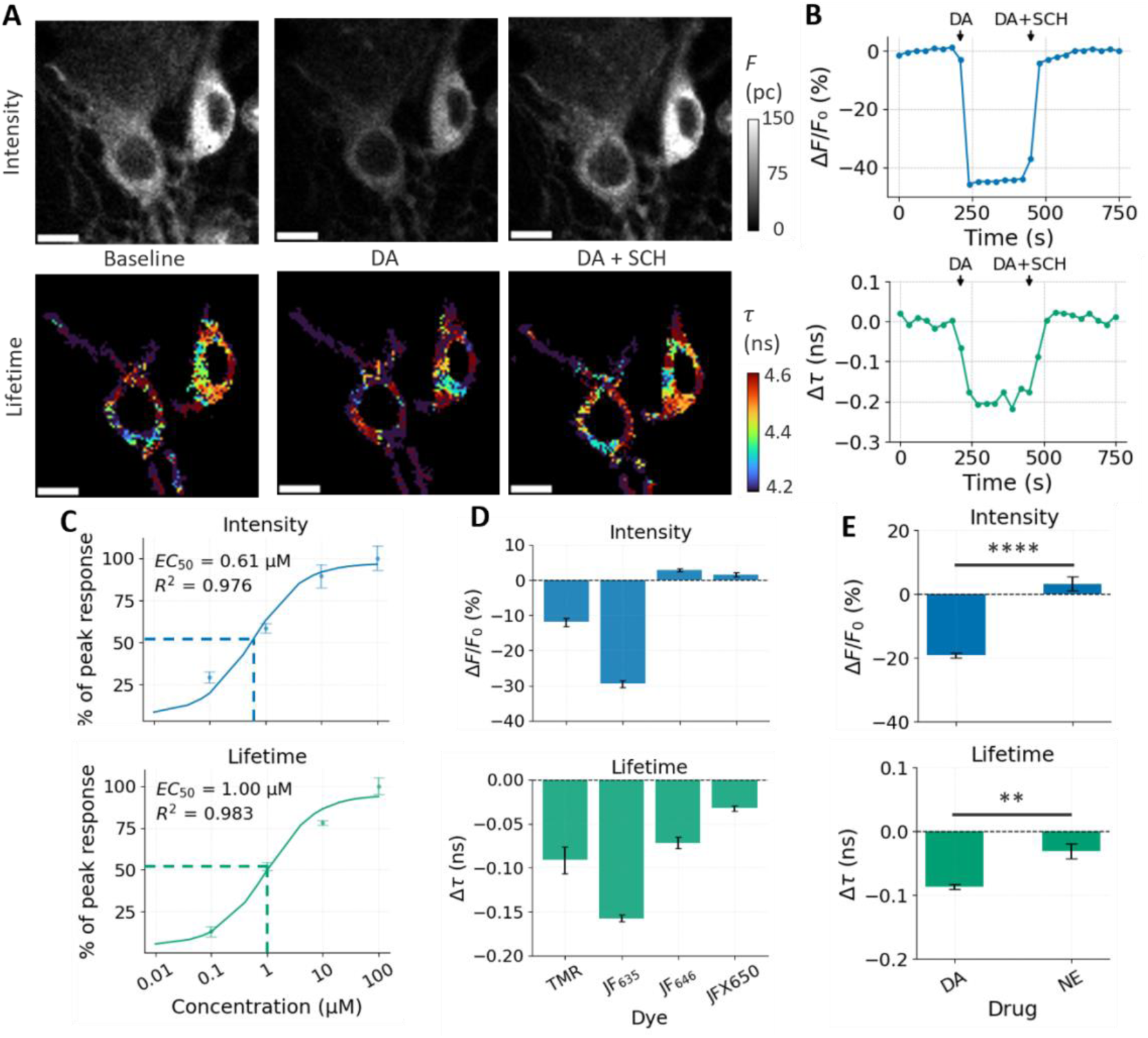
Pharmacological characterization and sensitivity of dHaloLife in primary neurons. **(A)** Representative intensity (top row) and fluorescence lifetime (bottom row) images of primary neurons expressing dHaloLife labeled with JF_635_. Cells were imaged at baseline, after addition of 100 µM dopamine (DA), and after 100 µM DA co-application with the DRD1-specific antagonist SCH (10 µM). Scale bars: 10 µm. Intensity was reported as photon counts (pc). **(B)** Representative time traces showing the antagonist-induced reversal of the sensor response in both fluorescence intensity (*ΔF/F_0_*, top) and lifetime (*τ*, bottom) upon sequential addition of DA and DA+SCH. Time traces represent the mean ± s.e.m. of the response for the highlighted cells in (A). **(C)** Dose-response titration of dHaloLife_635_. The sensor exhibited a sigmoidal response to DA with EC_50_ of 610 nM and 1,000 nM for the intensity and lifetime readouts, respectively. Data represent the mean ± s.e.m. from n ≥ 6 cells across ≥ 3 independent experiments. **(D)** Screening of dHaloLife against TMR, JF_635_, JF_646_, and JFX_650_. Data represent the mean ± s.e.m. from n ≥ 5 cells across ≥ 3 independent experiments. **(E)** Selectivity of dHaloLife_635_ against structurally related neurochemicals at 1 µM. Significant responses were observed only for DA, with marginal cross-reactivity to norepinephrine (NE). ****p < 0.0001, **p < 0.01, and n.s. (not significant) for p ≥ 0.05, two-tailed Welch’s t-test. Data represent the mean ± s.e.m. from n ≥ 6 cells across ≥ 3 independent experiments.

We next performed dose-response titrations in primary neurons across DA concentrations from 10 nM to 100 µM (**Figure 6C**). As in HEK293T cells, dHaloLife_635_ displayed a dose-dependent decrease in both intensity and lifetime (EC_50_ = 610 nM and 1,000 nM for the intensity and lifetime readouts, respectively, **Figure 6C**). Neurons thus showed a right-shifted response compared with HEK293T cells (EC_50_ = 190 nM and 880 nM, **Figure 3C**). Importantly, the photophysical distinction observed in HEK293T cells persisted: the intensity readout reached its half-maximal response at a lower DA concentration than the lifetime readout, consistent with the two readouts operating through distinct photophysical mechanisms.

To investigate whether the fluorophore-dependent lifetime modulation seen in our HEK293T matrix screen translated to primary neurons, we separately tested dHaloLife paired with TMR, JF_646_, and JFX_650_ in the presence of 100 µM DA (**Figure 6D**). In agreement with the HEK293T screen (**Figure 2B**), dHaloLife_635_ yielded the largest dynamic range in both intensity (*ΔF/F_0_* = -30.5 ± 2.8%) and lifetime (*Δτ* = -0.167 ± 0.007 ns, **Figure 6D**). With TMR, the sensor gave attenuated responses (*ΔF/F_0_* = -12.0 ± 2.8%, *Δτ* = -0.092 ± 0.034 ns). Conjugation with JF_646_ and JFX_650_ also mirrored the performance observed in HEK293T cells (**Figure 6D**), confirming that the cpHaloTag-dye pairings identified in HEK293T cells reliably predict sensor performance in primary neuronal cultures.

Finally, we tested the pharmacological selectivity of dHaloLife_635_ in primary neurons against norepinephrine (NE) and DA (1 µM, **Figure 6E**). Consistent with our HEK293T results (**Figure 3D**), dHaloLife_635_ demonstrated robust selectivity for DA (*ΔF/F_0_* = -19.1 ± 0.9%, *Δτ* = -0.086 ± 0.004 ns) over NE, which produced only a marginal response (*ΔF/F_0_* = 3.2 ± 2.2%, *Δτ* = -0.031 ± 0.011 ns, ****p < 0.0001 for intensity and **p < 0.01 for lifetime, two-tailed Welch’s t-test, **Figure 6E**). Together, these results demonstrated that dHaloLife_635_ preserves its high specificity, robust dynamic range, and quantitative lifetime stability in primary neurons, establishing its utility for functional neurochemical imaging.

## DISCUSSION

Fluorescence lifetime imaging microscopy (FLIM) has become increasingly important in biological studies because lifetime readouts are largely immune to intensity-related artifacts^20,21,43–45^. In biosensing applications spanning voltage^25^, metabolite^19^, second messenger^27,46^, and neurochemical^24,33^ sensing, lifetime readouts substantially improve measurement reliability and quantitative accuracy over intensity-based approaches^19,25^. The benefits extend beyond single-target sensing – combining FLIM with spectral imaging^47^ and pulsed interleaved excitation^48^ (PIE) has enabled highly multiplexed, concurrent imaging of multiple biological targets, reaching 6-plex imaging in live samples^22^ and 9-plex imaging in fixed samples^49^. Although the point-scanning requirement of conventional FLIM limits throughput, recent advances in electro-optic FLIM^25^ (EO-FLIM), a wide-field, all-optical approach to lifetime imaging, have increased imaging speeds by 2-3 orders of magnitude^44^. Together, these advances in FLIM instrumentation raise a fundamental question: can we develop a broader range of lifetime-based labels and sensors to match the growing capabilities and use of FLIM?

By mutating residues close to the chromophore’s electron-donating groups, Zhang’s group has developed a series of fluorescent proteins (FPs) that cover the visible spectrum (383-627 nm) and a wide range of lifetimes (1-5 ns)^50^. Similarly, Baker’s group used de novo protein design to develop NovoTags, which selectively bind green, orange, and far-red dyes with nanomolar affinity and exhibit tunable fluorescence lifetimes, enabling multiplexed imaging based on both emission wavelength and lifetime^51^. Using knowledge gained from crystal structures, molecular dynamics simulations and high-throughput screening, we have also developed a series of fluorogenic aptamers (e.g., Lettuce variants) that exhibit a range of lifetimes (from 4.4 to 6.0 ns) when binding their cognate small-molecule ligands^52,53^. However, both fluorescent proteins and fluorogenic aptamers are limited by their brightness and photostability. Johnsson’s group demonstrated HaloTag engineering as an effective means of diversifying lifetime-based chemigenetic labels^42^, which not only improves imaging quality but also achieves a large tuning range that facilitates multiplexed imaging. For instance, with engineered HaloTags, this tuning range can be as large as 2.89 ns, the difference between HaloTag9-JF_614_ (3.94 ns) and HaloTag10-JF_614_ (1.05 ns)^42^. Three of their variants, HaloTag7, HaloTag9, and HaloTag10, were also explored here (see **Supplementary Table 2**).

Tuning the lifetime of a label, however, is a different problem from making that lifetime respond to analyte binding. For instance, although the HaloTag-based chemigenetic calcium sensor HaloCaMP1b-JF_635_, which relies on an L-Z equilibrium shift, shows a +920% fluorescence intensity change upon Ca^2+^ binding^32^, no lifetime change has been reported for it. Only more recently have researchers reported a modified chemigenetic calcium sensor, WHaloCaMP1a_669_, with a +2.1 ns lifetime change (*Δτ* ) upon Ca^2+^ binding^27^ and a force sensor, WHaloForce-JFX_554_, with a *Δτ* of +0.02-0.04 ns in cardiomyocyte beating experiments^54^. Notably, the sensing mechanism of the WHalo sensors^27,54^, a shift in photoinduced electron transfer (PET), differs from that of HaloCaMP^32^, a shift in L-Z equilibrium. We will also explore this PET shift strategy in our future dHaloLife sensor designs.

The same difficulty applies to efforts to convert cpFP-based intensiometric sensors into lifetime-based sensors, such as GRAB_ACh3.0_ for acetylcholine (ACh) sensing^24^ and dLight3.8 for dopamine (DA) sensing^23^. Notably, Chen’s group tested several intensiometric GPCR sensor constructs (including GRAB_ACh3.0_, iAChSnFR, gGRAB_5-HT2h_, GRAB_NE2m_, and GRAB_DA2m_) and identified GRAB_ACh3.0_ as the construct with the largest lifetime change (*Δτ* = +0.17 ns) upon analyte binding^24^. Indeed, many cpFP-based sensors operate via equilibrium shifts that change absorbance rather than quantum yield, leading to strong intensity contrast with minimal lifetime response^18,54^. Our own screening bears this out: of 42 sensor variants tested, only dHaloLife_635_ showed a lifetime change exceeding 0.2 ns (*Δτ* = -0.21 ns) upon DA binding, underscoring the difficulty of obtaining a GPCR-based sensor with an appreciable lifetime response to analyte binding.

One of the most recent dLight sensors, dLight3.8, developed using knowledge gained from dLight1.3b and GRAB_ACh3.0_, shows the largest dynamic range (*∆F*/*F_0_*= 44) and permits robust, single-trial recording of DA release across a wide concentration range (1 nM to 100 μM) in response to electrical, optogenetic, and behavioral stimuli in multiple species and circuits^33^; however, its *Δτ* is only +0.24 ns upon DA binding. Compared with calcium sensors, which can exhibit a *Δτ* as large as +2.1 ns^27^, GPCR-based sensors have proven more challenging to engineer with an absolute *Δτ* exceeding 0.20 ns. Magnitude alone, however, is not the right figure of merit. Under shot-noise-limited conditions, a small *Δτ* in a bright sensor may provide more information than a large *Δτ* in a strongly quenched, dim sensor, because lifetime estimation depends on photon-counting statistics and strong quenching compromises measurement precision by reducing the available photon counts, despite the large lifetime dynamic range. Maintaining sufficient baseline brightness is therefore as essential for engineering lifetime sensors as it is for intensiometric sensors.

In this respect, the “turn-off” behavior of dHaloLife_635_, with a decrease in both fluorescence intensity and lifetime upon DA binding, may be an asset rather than a liability. Accurate lifetime measurement inherently relies on accumulating a sufficient photon budget across all functional states. In traditional “turn-on” intensiometric biosensors, such as HaloDA1.0^30^, a large increase in fluorescence intensity upon stimulation can introduce substantial dead-time counting losses that distort the recorded decay curves, or saturate highly sensitive single-photon detectors, such as avalanche photodiodes or photomultiplier tubes. By contrast, maintaining a bright baseline state ensures an adequate photon count even before stimulation, while a moderate intensity change upon analyte binding avoids detector saturation. An ideal lifetime sensor design may therefore favor a small intensity response paired with a large lifetime shift, balancing photon budget constraints against robust signal modulation.

It was previously reported that the JF_635_-HTL conjugate shows almost no visible absorption in aqueous solution (existing predominantly in the L form), but that absorbance and fluorescence increase more than 100-fold upon binding to the HaloTag^32^ (a shift to the Z form). However, HaloTag binding does not fully shift the dye’s equilibrium to the Z form, as the extinction coefficient of the JF_635_-HaloTag conjugate (*ε* = 81,000 M^−1^ cm^−1^) remains lower than the maximal value of free JF_635_ determined in acidic solution^38^ (*ε_max_* = 167,000 M^−1^ cm^−1^). This corresponds to an on-HaloTag *K_L-Z_* of 0.25 for JF_635_, suggesting that the photophysics of this far-red dye can be readily modulated by conformational changes within the HaloTag that alter its environment^32^. It has also been reported that in HEK293T cells, cpHaloTag-dye conjugates show fluorescence lifetimes comparable to those of HaloTag-dye conjugates, but with reduced brightness^54^. We focused on JF_635_ for our sensor constructs because it combines a large lifetime dynamic range (*Δτ* = -0.21 ns) with only a modest brightness reduction. Beyond this favorable trade-off, far-red chemigenetic constructs offer two further advantages over traditional cpFP-based sensors. First, HaloTag-dye conjugates are substantially brighter than common fluorescent proteins. HaloTag7 conjugated with JF_635_ is 3.4-fold and 7.1-fold brighter than EGFP and mCherry, respectively^55^ (**Supplementary Table 5**). Second, the HaloTag scaffold bypasses a major limitation of all FP-based sensors – their requirement for oxygen during chromophore maturation^56,57^, making dHaloLife_635_ a promising candidate for imaging DA in hypoxic environments where the performance of traditional cpFP-based sensors may be compromised^58^. We note, however, a recent trend toward replacing JF_635_-based sensors^32^ with those based on JF_525_^34^, JF ^30^, and JF ^27^, which offer improved bioavailability in the central nervous system^30^. Future work will focus on chemigenetic lifetime sensors with improved bioavailability.

Fluorescence lifetime is sensitive to local environmental conditions, including temperature, pH, ionic strength, and refractive index. Consequently, a calibration curve obtained using purified sensors in buffer at room temperature may not accurately reflect sensor responses in mammalian cells at 37 °C. In our experiments, dHaloLife_635_ exhibited slightly different baseline lifetimes (*τ_0_*) in HEK293T cells and primary neurons (4.59 ns and 4.56 ns, respectively), whereas the lifetime changes (*Δτ*) were of comparable magnitude (−0.212 ns and −0.167 ns, respectively). Similar baseline variations across expression systems have been documented previously for cpFP- and FRET-based lifetime sensors^24^. Because interconversion between bright and dark states and the associated photophysics can be complex^18^, the same sensor can exhibit different apparent EC_50_ values depending on the measurement modality. For DA sensing with dHaloLife_635_ in HEK293T cells, the EC_50_ values were 190 and 880 nM for intensity- and lifetime-based measurements, respectively. In primary neurons, the corresponding values were 610 and 1,000 nM. These modality- and cell-type-dependent differences underscore the need for calibration in the system in which the sensor will be used.

Optimizing lifetime-responsive sensors requires a different strategy from that used for traditional intensiometric sensors. Future optimization may be accelerated by computational structure prediction tools^59^, which could guide the rational design of the cpHaloTag-ligand interface to maximize environmental sensitivity. In addition, while traditional lifetime screening remains relatively low-throughput due to the need for point scanning, emerging lifetime-based flow cytometry platforms^60^ could enable high-throughput screening of large mutant libraries, substantially accelerating the optimization of next-generation sensors. To further improve the dynamic range (*Δτ*) of GPCR-based chemigenetic sensors, three complementary approaches will be needed: (1) structural insights from X-ray crystallography^50,52,53,61^, (2) dynamic information from high-resolution QM/MM (quantum mechanics/molecular mechanics) simulations, time-dependent density functional theory (TD-DFT)/MM simulations, and molecular dynamics simulations^50,62^, and (3) excited-state processes characterized by transient absorption measurements^50^. Together, these will guide the engineering of nonradiative (*k_nr_*) and radiative (*k_r_*) decay pathways^61^ and the validation of lifetime sensor mechanisms^50^.

The scaffold itself can be broadened as well. Among the self-labeling protein tags developed to date^63^, only HaloTag has been widely engineered into circularly permuted forms for sensing applications, yielding far-red chemigenetic sensors for calcium^32,64^, voltage^32^, protein-protein interactions^64^, force^54^, hydrogen peroxide^59^ and dopamine^30^. Although cpSNAP-tag and cpFAST have also been used to create calcium^65^ and hydrogen peroxide^66^ sensors, respectively, such examples remain rare. If additional self-labeling proteins could be converted into sensor scaffolds, the resulting orthogonal chemigenetic sensors would enable highly specific, multiplexed sensing.

To the best of our knowledge, dHaloLife_635_ is the first far-red-emitting, lifetime-responsive chemigenetic DA sensor built on a GPCR-cpHaloTag scaffold, distinguishing it from the intensity-based chemigenetic DA sensor HaloDA1.0^30^ and the cpFP-based DA sensor dLight3.8^33^. It arrives as the limitations of intensiometric sensors become more widely recognized and the field shifts toward sensors that exploit intrinsic photophysical properties such as lifetime (**Supplementary Table 9**), part of a broader push in neuroengineering toward quantitative, reproducible, and reliable readouts. Realizing that potential will require larger lifetime changes for improved signal-to-noise, which in turn depends on elucidating how the quantum yields of conjugated dyes are modulated within the HaloTag platform. dHaloLife_635_ is the first step toward that goal.

## ACKNOWLEDGEMENT

We thank Dr. Y. Li, Y. Chen and P. Ma for sharing their pdisplay-CMV-GRAB_ACh3.0_ and pdisplay-GRAB_ACh3.0mut_ constructs to test the imaging system in the Yeh lab. H.-C.Y. was supported by the National Science Foundation grants (CBET2432379 and CHE2404334 to H.-C.Y.) and the National Institutes of Health grants (DA060543 to H.-C.Y. and S.K.S. and UH3TR003148 to S.K.S). L.E.F. is supported by the Clayton Foundation for Research, W.M. Keck Foundation, BD2 Foundation, WoodNext Foundation, the National Science Foundation (CBET2432379), and the Waggoner Center for Alcohol and Addiction Research.

## AUTHOR CONTRIBUTIONS

A.-T.N., L.C. and H.-C.Y. discussed and defined the project. A.-T.N. and L.C. designed the sensors. A.-T.N., H.T. and Y.Z. cloned the constructs, performed FLIM measurements, including screening and characterization of the sensors, analyzed and organized the data. A.C., S.B. and L.E.F. produced the virus and prepared primary hippocampal neurons. A.-T.N. developed Python scripts for data analysis and illustration. Y.H., S.K., Y.-I.C., T.D.N., S.H., Y.-A.K., S.S. and W.-R.C. provided critical experimental support and insights and contributed to discussion. A.-T.N. and H.-C.Y. wrote the article with editorial assistance from all co-authors. H.-C.Y., S.K.S. and L.E.F. supervised the project.

## COMPETING INTERESTS

L.E.F. is a founder and officer of NumbCorp, on the scientific advisory board of Modulight.bio and receives consulting income for work related to Parkinson’s disease, optogenetics, cellular biologics and brain-computer interfaces.

## DATA AND MATERIALS AVAILABILITY

All data needed to evaluate the conclusions in the paper are present in the paper and/or the Supplementary Materials. Other data are available from the corresponding author upon reasonable request.

## Notes

### Competing Interest Statement

The authors have declared no competing interest.

### Summary of Updates

Results, methods, and SI were updated.

## REFERENCES

(1) Y. Wang, M. Wang, S. Yin, R. Jang, J. Wang, Z. Xue, T. Xu, “NeuroPep: a comprehensive resource of neuropeptides,” Database, 2015, bav038, 2015.

(2) Z. Wu, D. Lin, Y. Li, “Pushing the frontiers: tools for monitoring neurotransmitters and neuromodulators,” Nature Reviews Neuroscience, 23, 257–274, 2022.

(3) C. Dong, Y. Zheng, K. Long-Iyer, E. C. Wright, Y. Li, L. Tian, “Fluorescence imaging of neural activity, neurochemical dynamics, and drug-specific receptor conformation with genetically encoded sensors,” Annual Review of Neuroscience, 45, 2022.

(4) B. L. Sabatini, L. Tian, “Imaging neurotransmitter and neuromodulator dynamics in vivo with genetically encoded indicators,” Neuron, 108, 17–32, 2020.

(5) F. Nadim, D. Bucher, “Neuromodulation of neurons and synapses,” Current opinion in neurobiology, 29, 48–56, 2014.

(6) L. A. Gunaydin, L. Grosenick, J. C. Finkelstein, I. V. Kauvar, L. E. Fenno, A. Adhikari, S. Lammel, J. J. Mirzabekov, R. D. Airan, K. A. Zalocusky, “Natural neural projection dynamics underlying social behavior,” Cell, 157, 1535–1551, 2014.

(7) T. Patriarchi, J. R. Cho, K. Merten, M. W. Howe, A. Marley, W.-H. Xiong, R. W. Folk, G. J. Broussard, R. Liang, M. J. Jang, H. Zhong, D. Dombeck, M. von Zastrow, A. Nimmerjahn, V. Gradinaru, J. T. Williams, L. Tian, “Ultrafast neuronal imaging of dopamine dynamics with designed genetically encoded sensors,” Science, 360, eaat4422, 2018.

(8) A. Manglik, T. H. Kim, M. Masureel, C. Altenbach, Z. Yang, D. Hilger, M. T. Lerch, T. S. Kobilka, F. S. Thian, W. L. Hubbell, “Structural insights into the dynamic process of β2-adrenergic receptor signaling,” Cell, 161, 1101–1111, 2015.

(9) D. Hilger, M. Masureel, B. K. Kobilka, “Structure and dynamics of GPCR signaling complexes,” Nature Structural & Molecular Biology, 25, 4–12, 2018.

(10) W. I. Weis, B. K. Kobilka, “The molecular basis of G protein–coupled receptor activation,” Annual Review of Biochemistry, 87, 897–919, 2018.

(11) F. Sun, J. Zeng, M. Jing, J. Zhou, J. Feng, S. F. Owen, Y. Luo, F. Li, H. Wang, T. Yamaguchi, Z. Yong, Y. Gao, W. Peng, L. Wang, S. Zhang, J. Du, D. Lin, M. Xu, A. C. Kreitzer, G. Cui, Y. Li, “A genetically encoded fluorescent sensor enables rapid and specific detection of dopamine in flies, fish, and mice,” Cell, 174, 481–496. e419, 2018.

(12) M. Jing, P. Zhang, G. Wang, J. Feng, L. Mesik, J. Zeng, H. Jiang, S. Wang, J. C. Looby, N. A. Guagliardo, “A genetically encoded fluorescent acetylcholine indicator for in vitro and in vivo studies,” Nature Biotechnology, 36, 726–737, 2018.

(13) M. Jing, Y. Li, J. Zeng, P. Huang, M. Skirzewski, O. Kljakic, W. Peng, T. Qian, K. Tan, J. Zou, S. Trinh, R. Wu, S. Zhang, S. Pan, S. A. Hires, M. Xu, H. Li, L. M. Saksida, V. F. Prado, T. J. Bussey, M. A. M. Prado, L. Chen, H. Cheng, Y. Li, “An optimized acetylcholine sensor for monitoring in vivo cholinergic activity,” Nature Methods, 17, 1139–1146, 2020.

(14) T. Patriarchi, A. Mohebi, J. Sun, A. Marley, R. Liang, C. Dong, K. Puhger, G. O. Mizuno, C. M. Davis, B. Wiltgen, M. v. Zastrow, J. D. Berke, L. Tian, “An expanded palette of dopamine sensors for multiplex imaging in vivo,” Nature Methods, 17, 1147–1155, 2020.

(15) J. Feng, C. Zhang, J. E. Lischinsky, M. Jing, J. Zhou, H. Wang, Y. Zhang, A. Dong, Z. Wu, H. Wu, W. Chen, P. Zhang, J. Zou, S. A. Hires, J. J. Zhu, G. Cui, D. Lin, J. Du, Y. Li, “A genetically encoded fluorescent sensor for rapid and specific in vivo detection of norepinephrine,” Neuron, 102, 745–761. e748, 2019.

(16) C. Dong, C. Ly, L. E. Dunlap, M. V. Vargas, J. Sun, I.-W. Hwang, A. Azinfar, W. C. Oh, W. C. Wetsel, D. E. Olson, L. Tian, “Psychedelic-inspired drug discovery using an engineered biosensor,” Cell, 184, 2779–2792. e2718, 2021.

(17) J. Wan, W. Peng, X. Li, T. Qian, K. Song, J. Zeng, F. Deng, S. Hao, J. Feng, P. Zhang, Y. Zhang, J. Zou, S. Pan, M. Shin, B. J. Venton, J. J. Zhu, M. Jing, M. Xu, Y. Li, “A genetically encoded sensor for measuring serotonin dynamics,” Nature Neuroscience, 24, 746–752, 2021.

(18) L. Ravotto, L. Duffet, X. Zhou, B. Weber, T. Patriarchi, “A bright and colorful future for G-protein coupled receptor sensors,” Frontiers in Cellular Neuroscience, 14, 67, 2020.

(19) Y.-I. Chen, Y.-J. Chang, S.-C. Liao, T. D. Nguyen, J. Yang, Y.-A. Kuo, S. Hong, Y.-L. Liu, H. G. Rylander, S. R. Santacruz, T. E. Yankeelov, H.-C. Yeh, “Generative adversarial network enables rapid and robust fluorescence lifetime image analysis in live cells,” Communications Biology, 5, 1–11, 2022.

(20) T. D. Nguyen, Y.-I. Chen, A.-T. Nguyen, S. Yonas, M. P. Sripati, Y.-A. Kuo, S. Hong, M. Litvinov, Y. He, H.-C. Yeh, “Two-photon autofluorescence lifetime assay of rabbit photoreceptors and retinal pigment epithelium during light-dark visual cycles in rabbit retina,” Biomedical Optics Express, 15, 3094–3111, 2024.

(21) Y. Sun, T. D. Nguyen, Y.-I. Chen, U. C. Coskun, S.-C. Liao, H.-C. Yeh In SPIE Proceedings - Multiphoton Microscopy in the Biomedical Sciences XXIII; SPIE: 2023; Vol. 12384, p 36–51.

(22) T. D. Nguyen, Y.-I. Chen, A. T. Nguyen, L. H. Chen, S. Yonas, M. Litvinlov, Y. He, Y.-A. Kuo, S. Hong, H. G. Rylander III, H.-C. Yeh, “Multiplexed imaging in live cells using pulsed interleaved excitation spectral FLIM,” Optics Express, 32, 3290–3307, 2024.

(23) B. Lodder, T. Kamath, E. Savenco, B. Röring, M. Siegel, J. A. Chouinard, S. J. Lee, C. Zagoren, P. Rosen, I. Hartman, J. Timmins, R. Adan, L. Tian, B. L. Sabatini, “Absolute measurement of fast and slow neuronal signals with fluorescence lifetime photometry at high temporal resolution,” Neuron, 113, 3554–3566. e3557, 2025.

(24) P. Ma, P. Chen, E. I. Tilden, S. Aggarwal, A. Oldenborg, Y. Chen, “Fast and slow: Recording neuromodulator dynamics across both transient and chronic time scales,” Science Advances, 10, eadi0643, 2024.

(25) A. J. Bowman, C. Huang, M. J. Schnitzer, M. A. Kasevich, “Wide-field fluorescence lifetime imaging of neuron spiking and subthreshold activity in vivo,” Science, 380, 1270–1275, 2023.

(26) Y.-I. Chen, Y.-J. Chang, T. D. Nguyen, C. Liu, S. Phillion, Y.-A. Kuo, H. T. Vu, A. Liu, Y.-L. Liu, S. Hong, P. Ren, T. E. Yankeelov, H.-C. Yeh, “Measuring DNA hybridization kinetics in live cells using a time-resolved 3D single-molecule tracking method,” Journal of the American Chemical Society, 2019.

(27) H. Farrants, Y. Shuai, W. C. Lemon, C. Monroy Hernandez, D. Zhang, S. Yang, R. Patel, G. Qiao, M. S. Frei, S. E. Plutkis, J. B. Grimm, T. L. Hanson, F. Tomaska, G. C. Turner, C. Stringer, P. J. Keller, A. G. Beyene, Y. Chen, Y. Liang, L. D. Lavis, E. R. Schreiter, “A modular chemigenetic calcium indicator for multiplexed in vivo functional imaging,” Nature Methods, 21, 1916–1925, 2024.

(28) D. M. Shcherbakova, “Near-infrared and far-red genetically encoded indicators of neuronal activity,” Journal of Neuroscience Methods, 362, 109314, 2021.

(29) D. M. Shcherbakova, O. V. Stepanenko, K. K. Turoverov, V. V. Verkhusha, “Near-infrared fluorescent proteins: multiplexing and optogenetics across scales,” Trends in Biotechnology, 36, 1230–1243, 2018.

(30) Y. Zheng, R. Cai, K. Wang, J. Zhang, Y. Zhuo, H. Dong, Y. Zhang, Y. Wang, F. Deng, E. Ji, Y. Cui, S. Fang, X. Zhang, H. Huang, K. Zhang, J. Wang, G. Li, X. Miao, Z. Wang, Y. Yang, S. Li, J. B. Grimm, K. Johnsson, E. R. Schreiter, L. D. Lavis, Z. Chen, Y. Mu, Y. Li, “In vivo multiplex imaging of dynamic neurochemical networks with designed far-red dopamine sensors,” Science, 388, eadt7705, 2025.

(31) L. D. Lavis, “Teaching old dyes new tricks: biological probes built from fluoresceins and rhodamines,” Annual Review of Biochemistry, 86, 825–843, 2017.

(32) C. Deo, A. S. Abdelfattah, H. K. Bhargava, A. J. Berro, N. Falco, H. Farrants, B. Moeyaert, M. Chupanova, L. D. Lavis, E. R. Schreiter, “The HaloTag as a general scaffold for far-red tunable chemigenetic indicators,” Nature Chemical Biology, 17, 718–723, 2021.

(33) J. I. Roshgadol, J. A. Chouinard, S. Majumder, E. C. Scott, K. Borges, K. M. Hagihara, N. Mancini, T. Steveson, T. Kamath, B. Lodder, R. Dalangin, N. Tjahjono, A. Pal, C. Soares-Cunha, P. R. Melugin, A. Marley, K. Mahe, K. Kurima, S. Takahashi, D. Nosaka, K. Murakami, L. A. Colgan, P. T. Freitas, R. Chaudhuri, C. A. Siciliano, A. J. Rodrigues, V. Gradinaru, M. Von Zastrow, K. Podgorski, B. L. Sabatini, S. S. Bidaye, T. D. Hanks, N. Ji, J. R. Wickens, H. K. Inagaki, L. Tian, “Sensitive dLight3 for imaging broad-spectrum dopamine events across brain regions,” Research Square, rs. 3. rs-7313638, 2025.

(34) A. S. Abdelfattah, T. Kawashima, A. Singh, O. Novak, H. Liu, Y. Shuai, Y.-C. Huang, L. Campagnola, S. C. Seeman, J. Yu, J. Zheng, J. B. Grimm, R. Patel, J. Friedrich, B. D. Mensh, L. Paninski, J. J. Macklin, G. J. Murphy, K. Podgorski, B.-J. Lin, T.-W. Chen, G. C. Turner, Z. Liu, M. Koyama, K. Svoboda, M. B. Ahrens, L. D. Lavis, E. R. Schreiter, “Bright and photostable chemigenetic indicators for extended in vivo voltage imaging,” Science, 365, 699–704, 2019.

(35) Y. Nasu, Y. Shen, L. Kramer, R. E. Campbell, “Structure-and mechanism-guided design of single fluorescent protein-based biosensors,” Nature Chemical Biology, 17, 509–518, 2021.

(36) L. Wang, M. Tran, E. D’Este, J. Roberti, B. Koch, L. Xue, K. Johnsson, “A general strategy to develop cell permeable and fluorogenic probes for multicolour nanoscopy,” Nature Chemistry, 12, 165–172, 2020.

(37) J. B. Grimm, A. N. Tkachuk, L. Xie, H. Choi, B. Mohar, N. Falco, K. Schaefer, R. Patel, Q. Zheng, Z. Liu, “A general method to optimize and functionalize red-shifted rhodamine dyes,” Nature Methods, 17, 815–821, 2020.

(38) J. B. Grimm, A. K. Muthusamy, Y. Liang, T. A. Brown, W. C. Lemon, R. Patel, R. Lu, J. J. Macklin, P. J. Keller, N. Ji, L. D. Lavis, “A general method to fine-tune fluorophores for live-cell and in vivo imaging,” Nature Methods, 14, 987–994, 2017.

(39) C. Arnesano, Y. Santoro, E. Gratton, “Digital parallel frequency-domain spectroscopy for tissue imaging,” Journal of biomedical optics, 17, 096014, 2012.

(40) P. Kumar, J. D. Vevea, A. N. Tkachuk, E. R. Chapman, K. R. Campbell, E. T. Watson, D. J. Solecki, L. D. Lavis, “Optimizing multifunctional fluorescent ligands for intracellular labeling,” Proceedings of the National Academy of Sciences of the United States of America, 122, e2510046122, 2026.

(41) T. Patriarchi, J. R. Cho, K. Merten, A. Marley, G. J. Broussard, R. Liang, J. Williams, A. Nimmerjahn, M. von Zastrow, V. Gradinaru, “Imaging neuromodulators with high spatiotemporal resolution using genetically encoded indicators,” Nature Protocols, 14, 3471–3505, 2019.

(42) M. S. Frei, M. Tarnawski, M. J. Roberti, B. Koch, J. Hiblot, K. Johnsson, “Engineered HaloTag variants for fluorescence lifetime multiplexing,” Nature Methods, 19, 65–70, 2022.

(43) Y.-I. Chen, Y.-J. Chang, Y. Sun, S.-C. Liao, S. R. Santacruz, H.-C. Yeh, “Spatial resolution enhancement in photon-starved STED imaging using deep learning-based fluorescence lifetime analysis,” Nanoscale, 15, 9449–9456, 2023.

(44) F. Lin, C. Zhang, Z. Huang, Y. Wang, M. Yi, J. Li, X. Weng, Y. Chen, P. Lai, J. Qu, “Advances in fluorescence lifetime imaging microscopy: Techniques and biomedical applications,” Applied Physics Reviews, 13, 2026.

(45) B. Torrado, B. Pannunzio, L. Malacrida, M. A. Digman, “Fluorescence lifetime imaging microscopy,” Nature Reviews Methods Primers, 4, 80, 2024.

(46) F. H. van der Linden, E. K. Mahlandt, J. J. Arts, J. Beumer, J. Puschhof, S. M. de Man, A. O. Chertkova, B. Ponsioen, H. Clevers, J. D. van Buul, M. Postma, T. W. J. Gadella, J. Goedhart, “A turquoise fluorescence lifetime-based biosensor for quantitative imaging of intracellular calcium,” Nature Communications, 12, 7159, 2021.

(47) L. Scipioni, A. Rossetta, G. Tedeschi, E. Gratton, “Phasor S-FLIM: a new paradigm for fast and robust spectral fluorescence lifetime imaging,” Nature Methods, 18, 542–550, 2021.

(48) B. K. Muller, E. Zaychikov, C. Brauchle, D. C. Lamb, “Pulsed interleaved excitation,” Biophysical Journal, 89, 3508–3522, 2005.

(49) T. Niehorster, A. Loschberger, I. Gregor, B. Kramer, H. J. Rahn, M. Patting, F. Koberling, J. Enderlein, M. Sauer, “Multi-target spectrally resolved fluorescence lifetime imaging microscopy,” Nature Methods, 2016.

(50) Z. Tan, C.-H. Hsiung, J. Feng, Y. Zhang, Y. Wan, J. Chen, K. Sun, P. Lu, J. Zang, W. Yang, Y. Gao, J. Yin, T. Zhu, Y. Lu, Z. Pan, Y. Zou, C. Liao, X. Li, Y. Ye, Y. Liu, X. Zhang, “Time-resolved fluorescent proteins expand fluorescent microscopy in temporal and spectral domains,” Cell, 188, 6987–7005. e6928, 2025.

(51) L. Tran, S. Klein, D. Juergens, S. Sharma, J. Decarreau, G. R. Lee, Y. Wang, W. Chen, A. K. Bera, A. Kang, J. Woods, E. Joyce, D. K. Vafeados, N. Roullier, X. Li, B. Liu, Y. Bo, E. Muratspahić, T. A. Brown, J. B. Grimm, R. Patel, L. D. Lavis, J. Mahamid, L. An, D. Baker, “De novo design of orthogonal far-red, orange, and green fluorophore-binding proteins for multiplexed imaging,” Science, 393, 813–819, 2026.

(52) Y.-A. Kuo, Y.-I. Chen, N. Siraj, Y. He, Z. Yang, Y. Wang, E. J. Batchelder-Schwab, Z. Korkmaz, S. Yonas, T. D. Nguyen, S. Hong, A.-T. Nguyen, S. Kim, S. Seifi, P.-H. Fan, Y. Wu, H.-W. Liu, Y. Lu, P. Ren, C. Mao, H.-C. Yeh, “Fluorogenic aptamer optimization on a massively parallel sequencing platform,” ACS Sensors, 2026.

(53) Y.-I. Chen, Y.-A. Kuo, Y. He, N. Siraj, E. J. Batchelder-Schwab, Y.-J. Chang, S. Yonas, Y. Wu, Z. Yang, A.-T. Nguyen, “Lifetime-based multiplexed detection of viral RNA using fluorogenic aptamers,” bioRxiv, 2026.

(54) D. Wu, E. A. Morales, J. Lee, S. Pangeni, H. Farrants, K. A. Hutchings, R. J. Barndt, Q. Yu, X. Li, H. Shroff, H. Wu, A. G. Tebo, L. D. Lavis, E. R. Schreiter, T. Ha, S. Wang, “WHaloForce enables chemigenetic imaging of molecular tension in living cells and animals,” bioRxiv, 2026.2008. 2018.745627, 2026.

(55) T. J. Lambert, “FPbase: a community-editable fluorescent protein database,” Nature Methods, 16, 277–278, 2019.

(56) R. Heim, D. C. Prasher, R. Y. Tsien, “Wavelength mutations and posttranslational autoxidation of green fluorescent protein,” Proceedings of the National Academy of Sciences, 91, 12501–12504, 1994.

(57) A. Cook, F. Walterspiel, C. Deo, “HaloTag-based reporters for fluorescence imaging and biosensing,” ChemBioChem, 24, e202300022, 2023.

(58) K. Charubin, H. Streett, E. T. Papoutsakis, “Development of strong anaerobic fluorescent reporters for Clostridium acetobutylicum and Clostridium ljungdahlii using HaloTag and SNAP-tag proteins,” Applied and Environmental Microbiology, 86, e01271–01220, 2020.

(59) J. D. Lee, A. Nguyen, C. E. Gibbs, Z. R. Jin, Y. Wang, A. Moghadasi, S. J. Wait, H. Choi, K. M. Evitts, A. Asencio, S. B. Bremner, S. Zuniga, V. Chavan, I. K. Pranoto, C. A. Williams, A. Smith, F. Moussavi-Harami, M. Regnier, D. Baker, J. E. Young, D. L. Mack, E. Nance, P. M. Boyle, A. Berndt, “Monitoring in real time and far-red imaging of H2O2 dynamics with subcellular resolution,” Nature Chemical Biology, 1–12, 2025.

(60) H. Kanno, K. Hiramatsu, H. Mikami, A. Nakayashiki, S. Yamashita, A. Nagai, K. Okabe, F. Li, F. Yin, K. Tominaga, “High-throughput fluorescence lifetime imaging flow cytometry,” Nature Communications, 15, 7376, 2024.

(61) S. Mukherjee, P. Manna, S.-T. Hung, F. Vietmeyer, P. Friis, A. E. Palmer, R. Jimenez, “Directed evolution of a bright variant of mCherry: suppression of nonradiative decay by fluorescence lifetime selections,” The Journal of Physical Chemistry B, 126, 4659–4668, 2022.

(62) Y. He, Z. Yang, Y.-A. Kuo, Y. Wu, D. Fonseca-Albert, K. K. Le, J. Guo, Y. Wang, A.-T. Nguyen, Y.-I. Chen, S. Kim, W.-R. Chen, S. Seifi, S. Hong, T. D. Nguyen, Y. Chen, P. Ren, Y. Lu, H.-C. Yeh, “High-throughput on-chip screening enables rapid adaptation of DNA aptamers to SARS-CoV-2 evolution,” ACS Nano, 2026.

(63) N. Porzberg, K. Gries, K. Johnsson, “Exploiting covalent chemical labeling with self-labeling proteins,” Annual Review of Biochemistry, 94, 29–58, 2025.

(64) M.-C. Huppertz, J. Wilhelm, V. Grenier, M. W. Schneider, T. Falt, N. Porzberg, D. Hausmann, D. C. Hoffmann, L. Hai, M. Tarnawski, G. Pino, K. Slanchev, I. Kolb, C. Acuna, L. M. Fenk, H. Baier, J. Hiblot, K. Johnsson, “Recording physiological history of cells with chemical labeling,” Science, 383, 890–897, 2024.

(65) D. Zhang, Z. Chen, Z. Du, B. Bao, N. Su, X. Chen, Y. Ge, Q. Lin, L. Yang, Y. Hua, S. W. Xin Hua, F. Zho, N. Li, R. Liu, L. Jiang, C. Bao, Y. Zhao, J. Loscalzo, Y. Yang, L. Zhu, “Design of a palette of SNAP-tag mimics of fluorescent proteins and their use as cell reporters,” Cell Discovery, 9, 56, 2023.

(66) E. S. Potekhina, D. I. Bass, D. Ezeriņa, D. D. Fleckenstein, A. S. Chebotarev, V. A. Sysoeva, D. I. Maltsev, V. V. Pak, A. A. Moshchenko, A. I. Sokolov, I. N. Myasnyanko, M. S. Baranov, A. B. Fedotov, A. M. Zheltikov, A. A. Lanin, J. Messens, V. V. Belousov, “A color-tailored fluorogenic sensor for hydrogen peroxide,” Nature Chemical Biology, 2025.

