## Supplementary Information for "A Chemigenetic Fluorescence Lifetime Biosensor for Dopamine Sensing"

**Table of contents**

**Methods**

**Supplementary Figures**

Supplementary Figure S1. Simple schematic of dHaloLife.

Supplementary Figure S2. Design, structural prediction, and primary amino acid sequence of the dHaloLife dopamine biosensor.

Supplementary Figure S3. Chemical structure of the dyes with HaloTag ligand used in this study.

Supplementary Figure S4. Principles of frequency-domain fluorescence lifetime phasor analysis.

Supplementary Figure S5. Baseline phasor distributions of dHaloLife labeled with different dyes.

Supplementary Figure S6. 3D predicted structure of dHaloLife.

Supplementary Figure S7. Representative fluorescence intensity and lifetime traces of $dHaloLife_{635}$ upon dopamine addition and antagonist co-application in HEK293T.

Supplementary Figure S8. Fluorescence intensity and lifetime of $dHaloLife_{635}$ upon dopamine addition and antagonist co-application.

Supplementary Figure S9. Representative fluorescence intensity and lifetime traces of dHaloLife-$mut_{635}$ upon dopamine addition.

Supplementary Figure S10. Relative intensity and lifetime changes of $dHaloLife_{635}$ upon DA addition are unaffected by excitation laser power.

Supplementary Figure S11. Representative fluorescence intensity and lifetime traces of $dHaloLife_{635}$ in primary neurons.

Supplementary Figure S12. Schematic of our FLIM system.

Supplementary Figure S13. Comparison of fluorescence lifetimes obtained by digital frequency domain FastFLIM and time-correlated single-photon counting (TCSPC).

Supplementary Figure S14. Our system was validated with a published acetylcholine sensor that exhibited lifetime increases upon ligand binding $GRAB_{ACh3.0}$ in HEK293T cells.

Supplementary Figure S15. Our system was validated with a published acetylcholine sensor that exhibited lifetime increases upon ligand binding $GRAB_{ACh3.0}$ in primary neurons.

**Supplementary Tables**

Supplementary Table 1. Photophysical properties of rhodamine-based dyes.

Supplementary Table 2. Mutations and linkers used for all variants in this study.

Supplementary Table 3. Mean and s.e.m. of fluorescence intensity and lifetime of each variant-dye combination.

Supplementary Table 4. 95% confidence interval of mean of each variant-dye combination.

Supplementary Table 5. Statistical analysis of the fluorescence intensity of dHaloLife$_{635}$ to co-application of antagonist SCH-23390 (Supplementary Figure S8).

Supplementary Table 6. Statistical analysis of the fluorescence lifetime of dHaloLife$_{635}$ to co-application of antagonist SCH-23390 (Supplementary Figure S8).

Supplementary Table 7. Statistical analysis of the fluorescence intensity response of dHaloLife$_{635}$ to structurally related neurochemicals in HEK293T cells (Figure 3D).

Supplementary Table 8. Statistical analysis of the fluorescence lifetime response of dHaloLife$_{635}$ to structurally related neurochemicals in HEK293T cells (Figure 3D).

Supplementary Table 9. GPCR-based sensors for neurotransmitters and neuromodulators that exhibit lifetime change.

Supplementary Table 10. Fluorescence intensity change, lifetime change, baseline lifetime, 95% confidence interval (CI) of intensity and lifetime changes of the GRAB$_{ACh3.0}$ using the digital frequency domain FLIM in Yeh Lab.

Supplementary Table 11. Brightness comparison of dHaloLife to common fluorescent proteins.

Supplementary Table 12. dHaloLife DNA sequence.

**Supplementary Videos**

Supplementary Video 1. Time-lapse fluorescence lifetime imaging and phasor analysis of dHaloLife$_{635}$ in HEK293T cells.

Supplementary Video 2. Time-lapse fluorescence lifetime imaging and phasor analysis of dHaloLife-mut$_{635}$ in HEK293T cells.

Supplementary Video 3. Time-lapse fluorescence lifetime imaging and phasor analysis of dHaloLife$_{635}$ in primary neurons.

### Methods

#### DNA plasmid preparation

PCRs were performed with Q5 High-Fidelity 2× Master Mix (M0492S, New England Biolabs) according to the manufacturer's instructions. Primers for PCR amplification of DNA fragments were ordered from Integrated DNA Technologies with 20-bp overlaps. PCR products were purified with the Monarch PCR & DNA Cleanup Kit (5 μg) (T0130S, New England Biolabs), then assembled by Gibson assembly with the NEBuilder HiFi DNA Assembly Master Mix (E2621S, New England Biolabs). DH5α competent cells (T3007, Zymo Research) were transformed with the assembled product. Plasmid DNA was then isolated using the QIAprep Spin Miniprep Kit (27104, Qiagen). Final DNA constructs were verified by Sanger sequencing or whole-plasmid nanopore sequencing (Eton Biosciences). pCMV-dLight1.1 (Addgene plasmid #111053; a gift from the Tian group) and pAAV-synapsin-HaloCaMP1a-EGFP (Addgene plasmid #138327, a gift from the Schreiter group) were obtained from Addgene. Full DNA and amino acid sequence of dHaloLife were summarized in **Supplementary Figure S2** and **Supplementary Table 6**.

#### HEK293T cell culture

Human embryonic kidney cells (HEK293T, CRL-3216, ATCC) were cultured in Dulbecco's Modified Eagle's Medium (DMEM, D6429, Sigma-Aldrich) containing 10% (v/v) fetal bovine serum (FBS, 35-010-CV, Corning) and 1% penicillin-streptomycin (P/S, 15140122, Gibco) at 37 °C in 5% $CO_2$. HEK293T cells were plated on 35 mm glass-bottom dishes (P35G-1.5-14-C, MatTek Corporation) coated with fibronectin (F1141-1MG, Sigma-Aldrich) one day before transfection. Transfection was performed at 70-90% confluency using a mixture of 5 μg polyethylenimine (PEI MAX, 24765-100, Kyfora Bio) and 1 μg plasmid DNA in Opti-MEM I reduced serum medium (31985-070, Gibco), incubated for 30 min before addition to the cells.

Janelia Fluor dyes functionalized with a chloroalkane handle (i.e., HaloTag ligand), including $JF_{525}$-HTL (8805/10U, Bio-Techne), $JF_{635}$-HTL (8808/10U, Bio-Techne), $JF_{646}$-HTL (8809/10U, Bio-Techne), and $JFX_{650}$-HTL (HT1070, Promega), were reconstituted in DMSO to a stock concentration of 1 mM. TMR-HTL (G8251, Promega) was supplied as 15 μl of a 5 mM solution. Stocks were aliquoted and stored at -20°C.

One day before imaging, the cells were incubated overnight with 200 nM of the HTL-functionalized dye, then washed once with culture medium before imaging. Cells were imaged in HEPES-buffered Tyrode's solution (127 mM NaCl, 5 mM KCl, 1.8 mM $NaH_2PO_4$, 5.5 mM glucose, 1 mM $CaCl_2$, 1 mM $MgCl_2$, and 25 mM HEPES at pH 7.4). Cells were imaged 48 to 72 hours after transfection.

#### Primary hippocampal neuron culture

Male and female P4 Sprague-Dawley rat pups were cryoanesthetized before brains were extracted and hippocampi isolated and collected in ice cold dissection solution (160.8 mM NaCl, 4.96 mM KCl, 1.1 mM $MgSO_4 \cdot 7H_2O$, 3.87 mM $CaCl_2$, 5.03 mM HEPES, and 5.55 mM glucose).

Hippocampi were then digested for 25 minutes at 37°C in digestion solution (10 mL of dissection solution with the addition of 100 units of papain (P4762, Sigma), 100 units DNase I (EN0521, Thermo Scientific), 100 μL of 50 mM EDTA, 20 μL of 1N NaOH, 41.25 μL of 400 mM cysteine (aliquots stored at -20°C), and 100 μL of 100 mM $CaCl_2$). Following digestion, liquid was aspirated and replaced with a quenching solution (10 mL of serum medium supplemented with 25 mg bovine serum albumin, A3311, Sigma-Aldrich) and 25 mg trypsin inhibitor from Glycine max (T6522, Sigma-Aldrich). Hippocampi were left in quenching solution for 2 min at room temperature before the solution was aspirated and replaced with serum medium (203 mL Minimum Essential Medium (11090081, Gibco), 10 mL fetal bovine serum (A5670801, Gibco), 0.2 mL Mito+ Serum Extender (355006, Corning), and 765 mg glucose). The hippocampi were then gently triturated and passed through a 40 μm cell strainer (352235, Corning). The strained suspension was centrifuged at room temperature for 5 min at 300x g, the supernatant discarded, and the pellet gently resuspended in serum medium. The cell suspension was plated at approximately 75,000 cells per 35 mm dish directly onto the 14 mm glass insert (P35G-1.5-14-C, MatTek) precoated with Matrigel (356230, Corning) diluted 1:30 in Neurobasal Medium (21103049, Gibco). After a 2-hour incubation at 37°C in a 5% $CO_2$ humidity-controlled incubator, 3 mL of pre-warmed culture medium (250 mL Neurobasal Medium (21103049, Gibco), 3.12 mL fetal bovine serum (A5670801, Gibco), 10 mL B-27 Supplement (17504044, Gibco), and 2.5 mL CTS GlutaMAX Supplement (A1286001, Gibco)) was gently added to each dish, and the dishes were returned to the incubator. 65 hours after plating, a glial growth inhibition mixture was added to a final concentration of 0.03 mg/mL fluorodeoxyuridine (FUDR, F10705, RPI) and 0.025 mg/mL uridine (U3003, Sigma-Aldrich). At 6 days *in vitro* (DIV), 1 μL containing $1.8*10^{10}$ viral genomes of AAV-hSyn-dHaloLife-WPRE (see below) were bath-applied to each dish. Cells were imaged on DIV 13-22.

**Virus production**

The viruses used were prepared in-house. All AAV was produced with a serotype DJ packaging vector with AAV2 ITRs. Production used standard triple transfection of HEK293T/17 cells and iodixanol gradient ultracentrifugation purification. Briefly, HEK293T/17 cells were grown on 9 x 15 cm plates (12-600-004, Fisher Scientific) until 70-80% confluency was reached. For AAV transfection, the following DNA mix was used per 3 x 15 cm plates: 50 µg of pAdDeltaF6 helper plasmid (Addgene plasmid #112867), 50 µg of pAAV-DJ Rep/Cap plasmid (VPK-420-DJ, Cell Biolabs), and 30 µg of payload plasmid. The DNA mix was combined with 1.5 mL of DMEM (100% DMEM) and 390 µl of PEI (260085, Kyfora Bio). This transfection mix was left at room temperature for 10-20 min and then added to 30 mL of pre-warmed complete DMEM (89% DMEM, 10% FBS, 1% P/S). 10 mL of this solution was then added to each plate and incubated for 72 hours at 37°C in 5% $CO_2$. Cells were then scraped off the plate with a cell lifter (MSPP-229306, Celltreat), collected in PBS, and then pelleted at 800 g for 10 min. Cell pellets were resuspended, combined, and centrifuged again at 800 g for 10 min. If needed, cell pellets were frozen at -80°C as a pause point, before thawing at 37°C for 2 min. Pellets were lysed in 8 mL buffer containing 150 mM NaCl and 50 mM Tris-HCl and underwent two freeze-thaw cycles before being incubated with a final concentration of 50 U/mL benzonase nuclease (E1014, Sigma-Aldrich) for 30 min at 37°C. Tubes were swirled by hand every 10 min during incubation. Debris was cleared by centrifugation at

3000x g at 4°C for 15 min, and the clear supernatant was filtered through a 0.45 µm PES filter (76479-020, VWR). The clarified supernatant was loaded on top of a discontinuous iodixanol (D1556, Sigma-Aldrich) gradient consisting of 15%, 25%, 40% and 60% iodixanol layers in a 35 mL Quick-Seal ultracentrifuge tube (03-989, Thermo Scientific). This was spun at 350,000x g for 90 min at 12°C in a Sorvall WX+ ultracentrifuge using maximum acceleration and deceleration. Viral particles (2.5-3 mL) were collected by withdrawing the clear 40% fraction with an 18-gauge needle. Before concentrating the recovered virus fraction, 15 mL Amicon Ultra Centrifugal Filters, 100-kDa molecular weight cut-off (UFC9100, Millipore Sigma), were incubated with 10 mL of 0.1% Pluronic F68 (ICN2750049, MP Biomedicals) in PBS for 10 min at room temperature. This was then removed and replaced with 15 mL of 0.01% Pluronic F68 and spun at 3,000 g for 5 min at 4°C. Flow-through was discarded, and 3 - 4 mL of 0.001% of Pluronic F68 with 200 mM NaCl was loaded and spun at 3,000 g for 5 min at 4°C. The viral fraction was then loaded with 3-4 mL of 0.001% Pluronic F68 with 200 mM NaCl and centrifuged at 3,000 g for 4 min at 4°C. Flow-through was discarded, and the sample was centrifuged in 2 min intervals under the same conditions until 500 µL of the sample remained. This retained sample was then filtered through a 0.5 mL Amicon Ultra Centrifugal Filter, 100-kDa molecular weight cut-off (UFC510024, Millipore Sigma), at 2,500 g for 3 min, and the flow-through discarded. This was centrifuged in 2 min increments until a final volume of 70 µL was reached. The filter insert was then inverted and placed into a fresh collection tube, which was centrifuged at 2,000 g for 2 min. Purified virus was aliquoted into 2 µL aliquots and stored at -80°C. Genomic titer was determined by qPCR. Viral dilutions were calculated based on original titers.

**Fluorescence lifetime imaging of cultured cells**

All FLIM experiments were carried out on the ISS Alba v5 laser scanning system attached to a Nikon TiU microscope equipped with a Nikon 60× NA 1.2 water immersion objective (CFI Plan Apochromat Lambda 60×C, Nikon, **Supplementary Figure S12**). The system includes an ASI XY automated stage with motorized Z control. The 488 nm and 635 nm excitation light was provided by diode lasers (ISS) triggered by a 40 MHz internal source. The laser repetition period was 25 ns, divided into 256-time channels. Emission light, after passing through a bandpass filter (535/50 nm for $JF_{525}$; 585/40 nm for TMR; and 731/130 nm for $JF_{635}$, $JF_{646}$, and $JFX_{650}$, AVR Optics), was collected by an avalanche photodiode (SPCM-AQR-15, PerkinElmer) and analyzed by a digital frequency-domain module (DFD, FastFLIM, ISS). A 200 µm pinhole was used. Each image frame comprised 256 × 256 pixels (100 × 100 µm FOV) with a pixel dwell time of 50 µs. Photons from 6-8 frames were pooled for intensity and lifetime calculation, giving a sampling rate of 0.04-0.05 Hz. To establish the correct scale for phasor analysis, the phasor plot coordinates were calibrated with standards of a known lifetime, which also accounted for the instrument response function (IRF) and background. Before data acquisition, the FastFLIM system was calibrated with fluorescein (4.0 ns in PBS, pH 7.4) for 488 nm and Atto633 (3.3 ns in deionized water) for 635 nm excitation (calibration reference values provided by ISS). Phasor analysis was performed in VistaVision (ISS). Intensity data were analyzed in ImageJ (Fiji) and reported as the fractional intensity change ($\Delta F/F_0$), where $\Delta F$ is the intensity change and $F_0$ is the mean intensity of the unbound sensor. Raw intensity data were reported as total photon counts (pc). For instance, if a pixel accumulates 10 pc over 6 stacked frames with a pixel dwell time of 50 µs, the pixel was illuminated for a total of 300 µs, corresponding to a photon count rate of 33.3 kHz. Our DFD-FLIM system was validated against time-correlated single-photon counting (TCSPC) FLIM and yielded comparable fluorescence lifetime measurements (see **Supplementary Figure S13**). Additionally, our DFD-FLIM was validated using $GRAB_{ACh3.0}$ in HEK293T cells and neuron cells (see **Supplementary Figures S14** and **S15**).

**Pharmacology**

Unless otherwise noted, all chemicals were applied by customized bath perfusion. The lifetime baseline was monitored until it had remained stable for at least 5 min before drug application. Final concentrations were as follows: dopamine hydrochloride (DA, 1 nM to 100 µM, A11136.06, Thermo Scientific), the DRD1-specific antagonist SCH-23390 (R(+)-SCH-23390 hydrochloride, 10 µM, D054, Sigma-Aldrich), L-(−)-norepinephrine (+)-bitartrate salt monohydrate (NE, 1 µM, A9512, Sigma-Aldrich), serotonin hydrochloride (5-HT, 1 µM, B21263.03, Thermo Scientific), 3,4-dihydroxy-L-phenylalanine (L-DOPA, 1 µM, D9628, Sigma-Aldrich), γ-aminobutyric acid (GABA, 1 µM, A5835, Sigma-Aldrich), and acetylcholine chloride (ACh, 100 µM, A6625, Sigma-Aldrich).

**Fluorescence lifetime analysis using the phasor approach**

To analyze the lifetime data, the phasor approach was employed[1], where each image pixel was mapped to its corresponding phasor coordinates (G and S) and displayed in a phasor plot, creating a phasor distribution. For cellular imaging, only healthy cells (judged by gradient contrast images) with membrane expressions were selected. Cells with round shape or sensor expression aggregates were excluded. The membrane of individual cells was selected as region of interest (ROI). To minimize the effect of movement artifact on intensity measurement, pixels with photon count below 10% of max intensity were excluded. Photons from effective pixels of a given ROI were pooled for use in the following equation ($\sum N_i$ , where $N_i$ represents the photon counts at the $i$-th pixel). For whole field-of-view-based FLIM analysis, pixels with more than 300 photons were excluded to avoid dead time artifact of the FLIM driver board. Photons from the rest of the pixels in the field-of-view were pooled for further analysis. The average photon count per pixel was used for intensity measurement. The average lifetime ($\tau$) of all photons in this ROI was calculated as follows:

$$\tau = \frac{\sum [N_i \cdot \tau_i]}{\sum N_i}$$

where $N_i$ represents the photon counts and $\tau_i$ represents the lifetime at the $i$-th pixel within the ROI, respectively. Phasor analysis generated a phase lifetime ($\tau_p$) and a magnitude lifetime ($\tau_m$) at each pixel. Here $\tau_p$ was used as $\tau_i$ to represent the lifetime at the $i$-th pixel.

**Determination of the optimal modulation frequency**

For a single-lifetime species, the phasor distribution shifts leftward along the semicircle as the modulation frequency ω increases, increasing the S/G ratio. In DFD-FLIM, the optimal modulation frequency $\omega$ is given by[2]:

$$\omega_{optimal}^2 = \frac{1+\sqrt{3}}{2\tau^2}$$

Because dHaloLife$_{635}$ lifetime readouts were around 4.5 ns, the corresponding optimal frequency was ~41 MHz; a laser repetition rate of 40 MHz was therefore used in this study.

**AlphaFold2 structure prediction**

Structural prediction was performed with ColabFoldv1.6.1[3,4] under default settings (AlphaFold2 with MMseqs2 for multiple sequence alignment). Structures were visualized in UCSF ChimeraX[5].

**Statistics and reproducibility**

All experiments were conducted with at least three independent biological replicates. Summary data are reported as the mean ± standard error of the mean (s.e.m.). Box plots were used where cell-to-cell distributions are shown (**Figure 5**), all other panels show mean ± s.e.m. Graphs were plotted in Python. Differences were assessed by two-tailed Student's t-test or one-way/ two-way ANOVA, with *$p < 0.05$, **$p < 0.01$, ***$p < 0.001$, ****$p < 0.0001$, and n.s. (not significant) for $p \geq 0.05$ using GraphPad Prism. Dose-response curves were fitted to the Hill equation using the HillFit library in Python developed by Freiburger and Imoto[6].

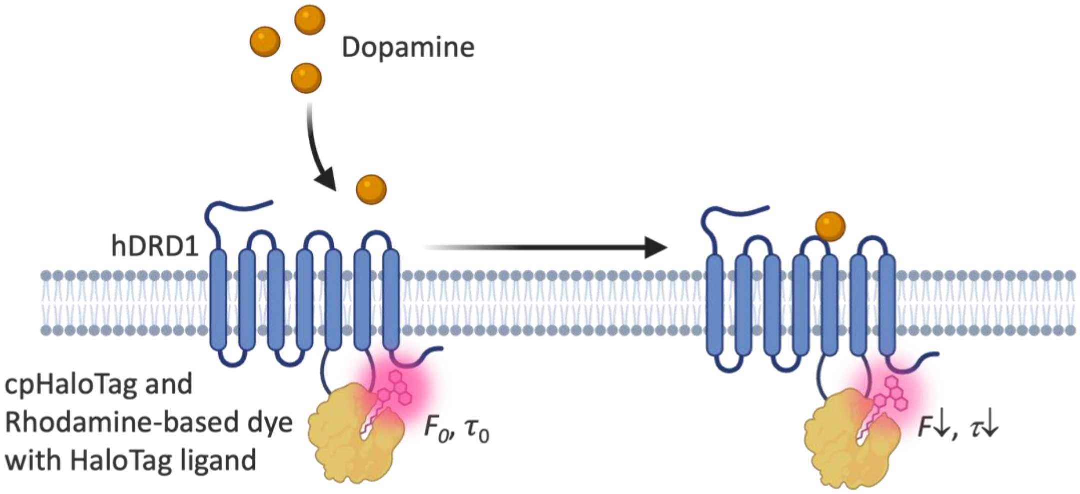


**Supplementary Figure S1. Simple schematic of dHaloLife.** Circularly permuted HaloTag (cpHaloTag) was inserted into the intracellular loop 3 of G-protein-coupled receptor (GPCR), specifically the human dopamine receptor D1 (hDRD1). The rhodamine-based dye with HaloTag ligand (HTL) was covalently bound to the cpHaloTag protein. Upon dopamine binding, the hDRD1 was activated and went through a conformational change, changing the local environment surrounding the HaloTag with dye, resulting in fluorescence intensity and lifetime changes. Schematics were made with BioRender. Figure adapted from Chen's group[7].

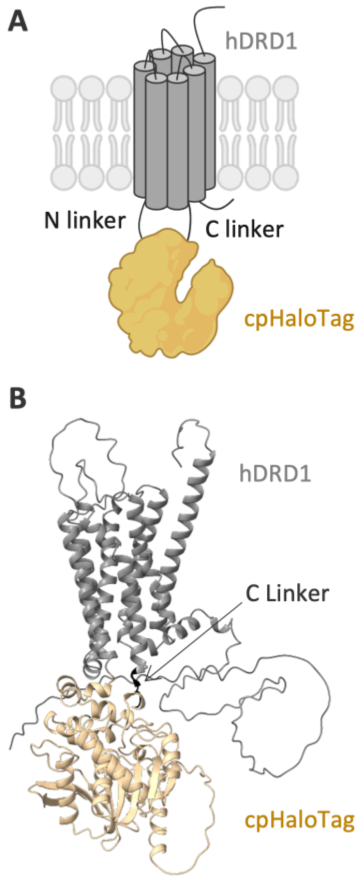


C

| | | |
|---|---|---|
| 1 | MRTLNTSAMDGTGLVVERDFSVRILTACFLSLLILSTLLG | 40 |
| 41 | NTLVCAAVIRFRHLRSKVTNFFVISLAVSDLLVAVLVMPW | 80 |
| 81 | KAVAEIAGFWPFGSFCNIWVAFDIMCSTASILNLCVISVD | 120 |
| 121 | RYWAISSPFRYERKMTPKAAFILISVAWTLSVLISFIPVQ | 160 |
| 161 | LSWHKAKPTSPSDGNATSLAETIDNCDSSLSRTYAISSSV | 200 |
| 201 | ISFYIPVAIMIVTYTRIYRIAQKETFQAFRTTDVGRKLII | 240 |
| 241 | DQNVFIEGTLPMGVVRPLTEVEMDHYREPFLNPVDREPLW | 280 |
| 281 | RFPNELPIAGEPANIVALVEEYMDWLHQSPVPKLLFWGTP | 320 |
| 321 | GVLIPPAEAARLAKSLPNCKAVDIGPGLNLLQEDNPDLIG | 360 |
| 361 | SEIARWLSTLEISGGGTGGSGGTGGSGGTGGSMAEIGTGF | 400 |
| 401 | PFDPHYVEVLGERMHYVDVGPRDGTPVLFLHGNPTSSYVW | 440 |
| 441 | RNIIPHVAPTHRCIAPDLIGMGKSDKPDLGYFFDDHVRFM | 480 |
| 481 | DAFIEALGLEEVVLVIHDWGSALGFHWAKRNPERVKGIAF | 520 |
| 521 | MEFIRPIPTWDEWPEFARKRETKVLKTLSVIMGVFVCCWL | 560 |
| 561 | PFFILNCILPFCGSGETQPFCIDSNTFDVFVWFGWANSSL | 600 |
| 601 | NPIIYAFNADFRKAFSTLLGCYRLCPATNNAIETVSINNN | 640 |
| 641 | GAAMFSSHHEPRGSISKECNLVYLIPHAVGSSEDLKKEEA | 680 |
| 681 | AGIARPLEKLSPALSVILDYDTDVSLEKIQPITQNGQHPT* | 720 |

**Supplementary Figure S2. Design, structural prediction, and primary amino acid sequence of the dHaloLife dopamine biosensor. (A)** Schematic representation of the general sensor design used in this study, featuring the human dopamine receptor D1 (hDRD1) fused to the circularly permuted HaloTag with N-terminal and C-terminal linkers. **(B)** Predicted structure of dHaloLife. Note that the dHaloLife design lacks an N-terminal linker. **(C)** Complete primary amino acid sequence of dHaloLife (720 amino acids). Color-coding designates the N-terminal domain of the hDRD1 receptor sequence (gray), followed by the C-terminal domain of the HaloTag (yellow), a flexible linker connecting the 2 domains of the HaloTag, then the N-terminal domain of the HaloTag (yellow), creating a cpHaloTag, then C-linker of the sensor (black) followed by the C-terminal domain of the hDRD1 (gray).

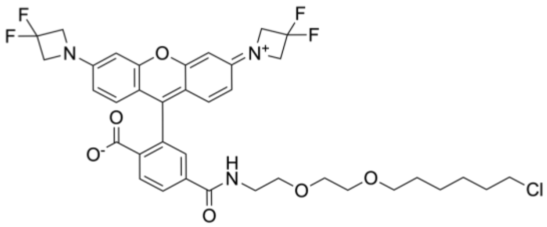

$JF_{525}$ – HaloTag ligand

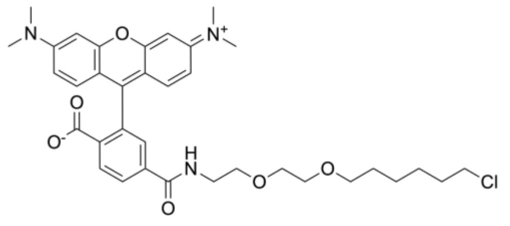

TMR – HaloTag ligand

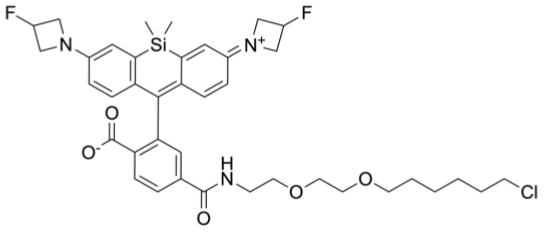

$JF_{635}$ – HaloTag ligand

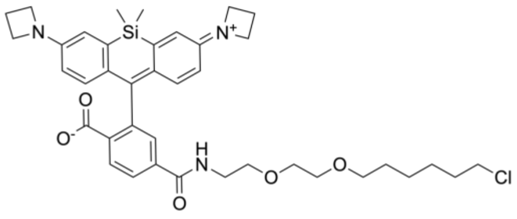

$JF_{646}$ – HaloTag ligand

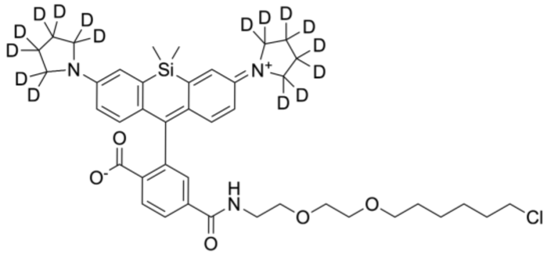

$JFX_{650}$ – HaloTag ligand

**Supplementary Figure S3. Chemical structure of the dyes with HaloTag ligand used in this study.**

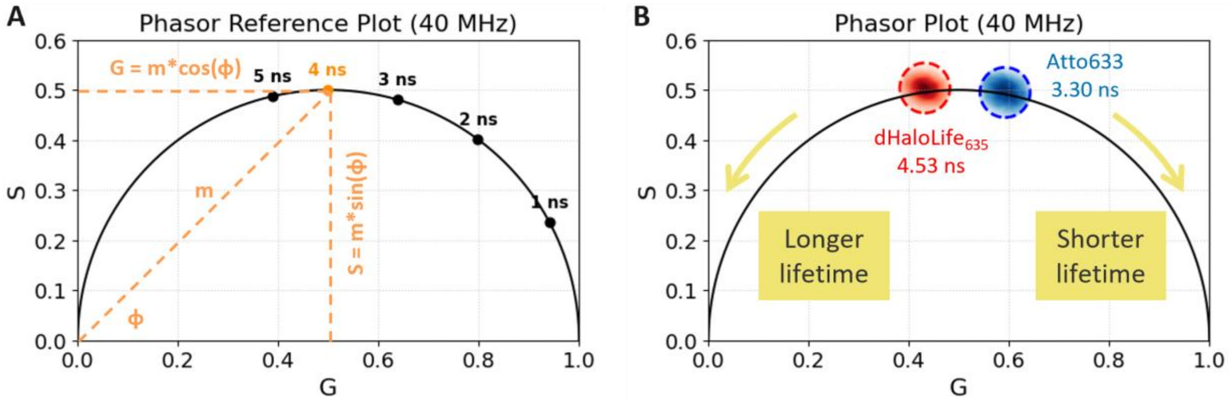


**Supplementary Figure S4. Principles of frequency-domain fluorescence lifetime phasor analysis. (A)** Phasor Reference Plot (40 MHz) illustrating the universal semicircle. The phasor approach transforms fluorescence lifetime data into a polar coordinate system plotted on a 2D Cartesian grid, where the horizontal axis represents the real component (G) and the vertical axis represents the imaginary component (S). For any given fluorescence lifetime (such as the highlighted 4 ns reference point in orange), the coordinates are mathematically defined by the sine and cosine of the phase shift angle (ϕ) and the modulation depth (m) relative to the modulated excitation source at 40 MHz. **(B)** Experimental Phasor Plot (40 MHz) of the baseline of dHaloLife$_{635}$ and the calibration dye Atto633. Experimental phasor distributions for specific labels mapped against the reference space. The plot displays the measured phasor cluster for the calibration dye Atto633 corresponding to a single-exponential lifetime of 3.30 ns (blue) and dHaloLife$_{635}$ corresponding to a single-exponential lifetime of 4.53 ns (red). All single-exponential fluorescence lifetimes lie on the boundary of the universal semicircle (diameter from G = 0 to 1, S = 0 to 0.5). Shorter lifetimes reside closer to (1, 0) (e.g., 1 ns) and longer lifetimes move counter-clockwise toward the origin (0, 0) (e.g., 5 ns).

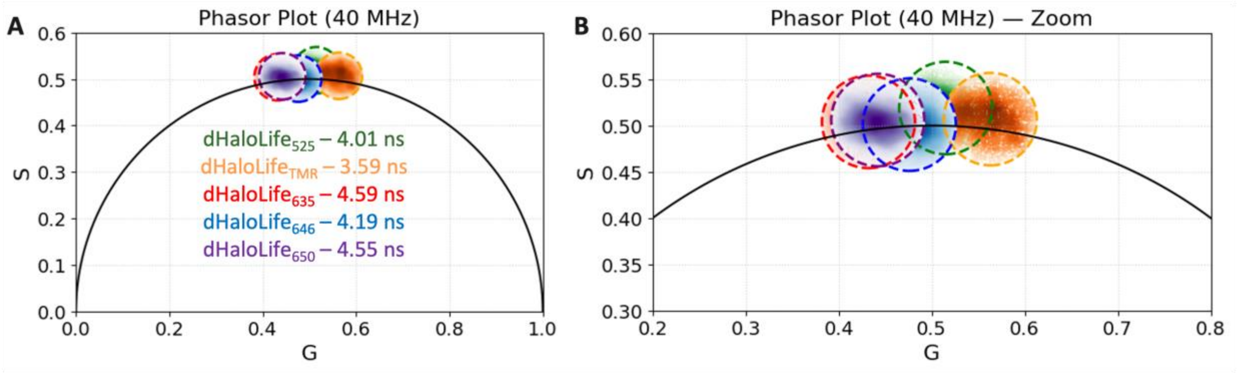


**Supplementary Figure S5. Baseline phasor distributions of dHaloLife labeled with different dyes. (A)** Representative phasor distributions at 40 MHz for dHaloLife labeled with $JF_{525}$ (green), TMR (orange), $JF_{635}$ (red), $JF_{646}$ (blue), and $JFX_{650}$ (purple). The corresponding lifetimes are indicated in the plot. **(B)** Enlarged view of the phasor distributions shown in **(A)**. The lifetime distributions lie on the universal semicircle black curve indicates lifetimes are single exponential.

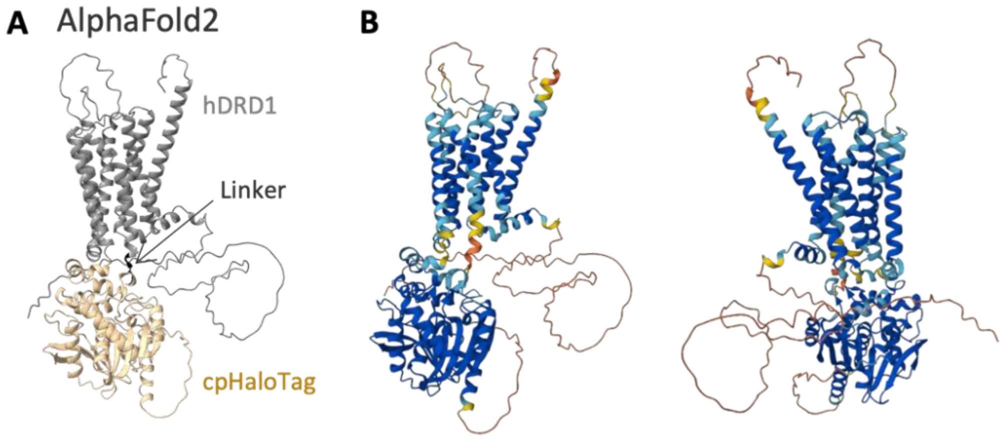


**Supplementary Figure S6. 3D predicted structure of dHaloLife. (A)** AlphaFold2 predicted structure (as seen in **Figure 2A**) with its predicted local distance difference test pLDDT score **(B)** (right sub-figure is the 180 degrees rotation of the left sub-figure). The average pLDDT score is 77.50 (confident).

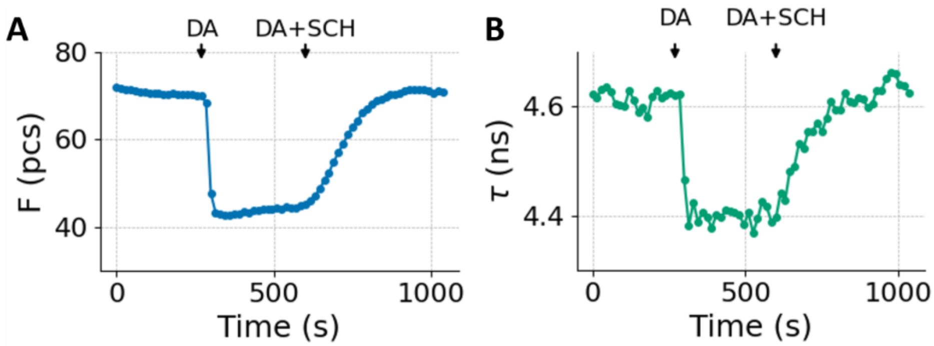


**Supplementary Figure S7. Representative fluorescence intensity and lifetime traces of dHaloLife$_{635}$ upon dopamine addition and antagonist co-application in HEK293T. (A)** Fluorescence intensity (*F*, photon counts) and **(B)** fluorescence lifetime ($\tau$, ns) of dHaloLife$_{635}$ over time. Arrows indicate dopamine (DA) addition and subsequent co-application of DA with the DRD1-specific antagonist SCH-23390 (SCH). Both intensity and lifetime decreased upon DA addition and recovered after antagonist co-application.

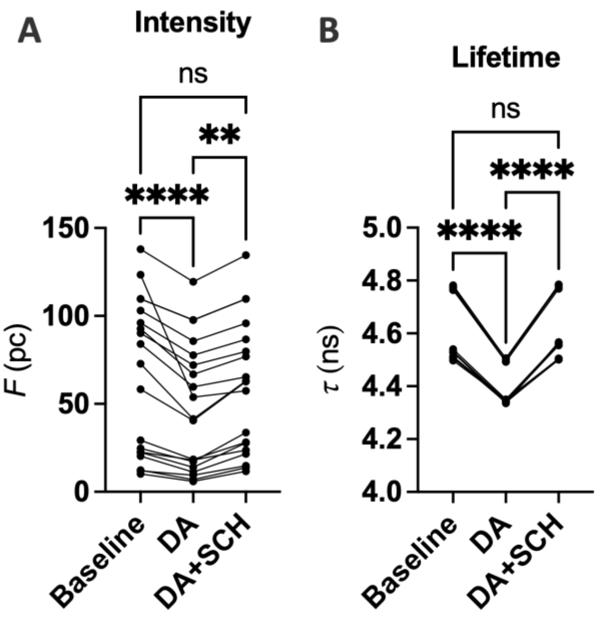


**Supplementary Figure S8. Fluorescence intensity and lifetime of dHaloLife$_{635}$ upon dopamine addition and antagonist co-application. (A)** Fluorescence intensity (*F*, photon counts) and **(B)** fluorescence lifetime ($\tau$, ns) at baseline, after DA addition, and after co-application of DA with SCH-23390 (DA+SCH). Each line connects measurements from the same cell (n = 19 cells from 3 independent experiments). Statistical results are reported in **Supplementary Tables 5** and **6**. n.s., not significant; **$p < 0.01$; ****$p < 0.0001$ using one-way repeated-measures ANOVA with Tukey's multiple comparisons test.

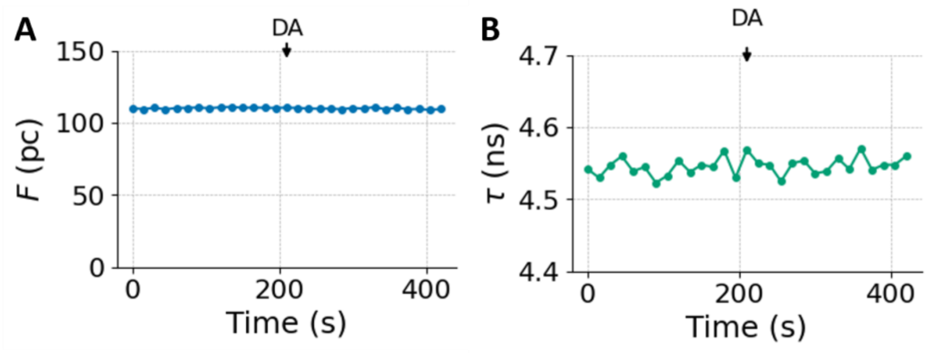


**Supplementary Figure S9. Representative fluorescence intensity and lifetime traces of dHaloLife-mut$_{635}$ upon dopamine addition. (A)** Fluorescence intensity (*F*, photon counts) and **(B)** fluorescence lifetime ($\tau$, ns) of dHaloLife-mut$_{635}$ over time. The arrow indicates DA addition. The traces show little change in fluorescence intensity or lifetime following DA addition.

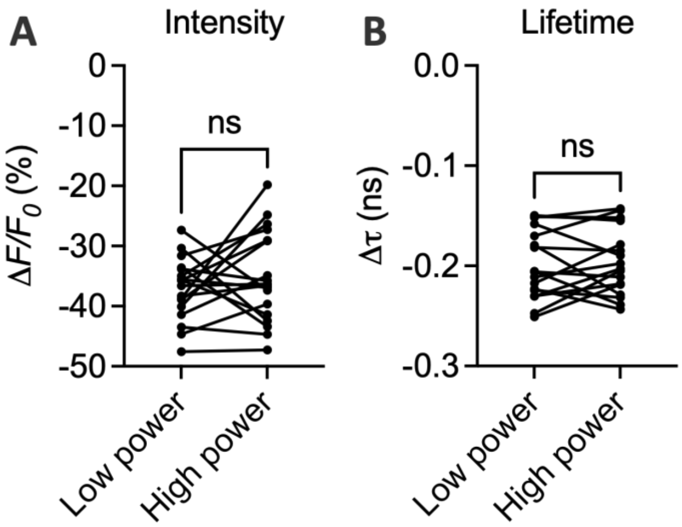


**Supplementary Figure S10. Relative intensity and lifetime changes of dHaloLife$_{635}$ upon DA addition are unaffected by excitation laser power. (A)** DA-induced change in fluorescence intensity (Δ$F/F_0$, %) and **(B)** DA-induced change in fluorescence lifetime (Δ$\tau$, ns) of dHaloLife635 in HEK293T cells measured at low and high excitation laser power. Each line connects the same cell imaged at both powers (n = 18 cells from 4 independent experiments). n.s. (not significant) using two-tailed paired Student's t-test.

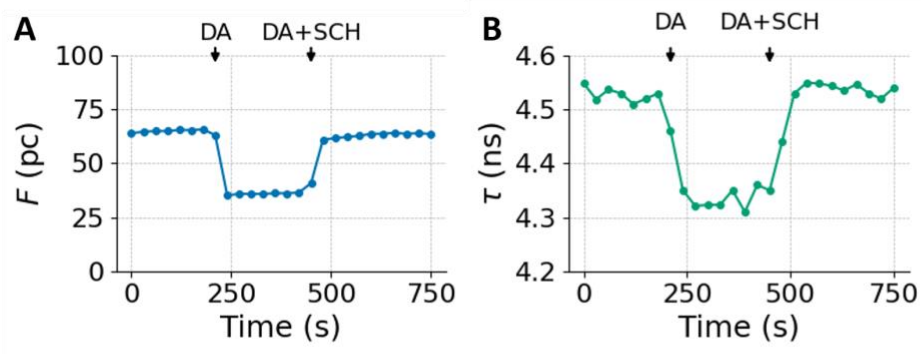


**Supplementary Figure S11. Representative fluorescence intensity and lifetime traces of dHaloLife$_{635}$ in primary neurons. (A)** Fluorescence intensity (*F*, photon counts) and **(B)** fluorescence lifetime ($\tau$, ns) of dHaloLife$_{635}$ in primary neurons over time. Arrows indicate DA addition and subsequent co-application of DA with SCH-23390 (DA+SCH). Both intensity and lifetime decreased upon DA addition and recovered after antagonist co-application.

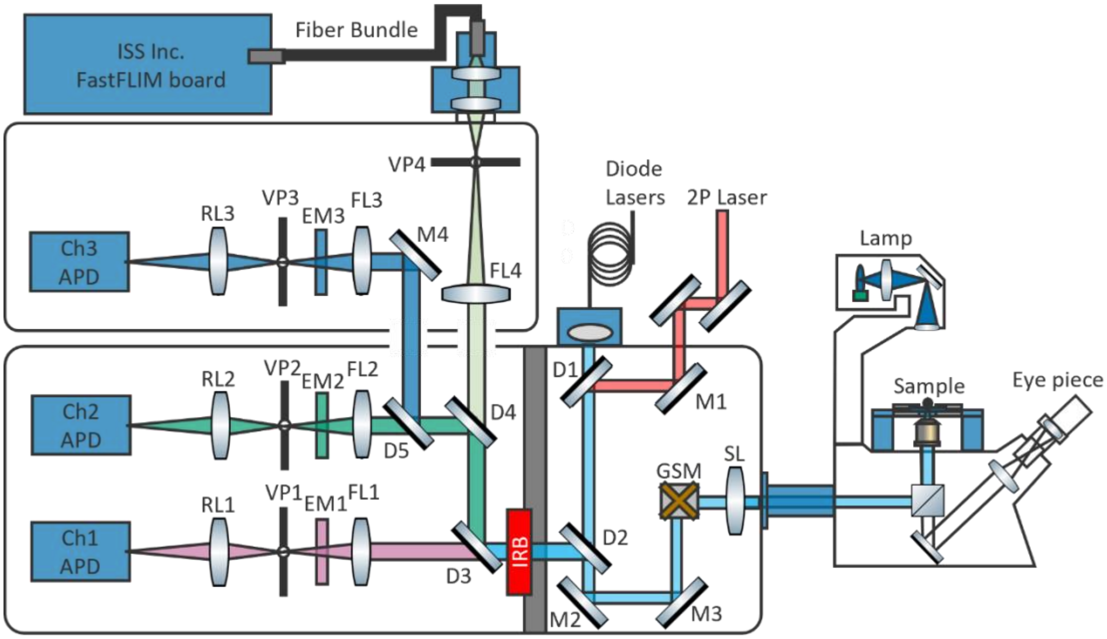


**Supplementary Figure S12. Schematic of our FLIM system.** Abbreviations: 2P – two-photon; APD – avalanche photodiode; RL – relay lens; VP – variable pinhole; EM – emission filter; FL – focusing lens; IRB – infrared blocking filter; M – mirror; D – dichroic mirror; GSM – galvo mirror; SL – scanning lens. All FLIM experiments were carried out on the ISS Alba v5 laser scanning system attached to a Nikon TiU microscope equipped with a Nikon 60× NA 1.2 water immersion objective (CFI Plan Apochromat Lambda 60×C, Nikon). The system includes an ASI XY automated stage with motorized Z control. The excitation light was provided by diode lasers (ISS) triggered by a 40 MHz internal source. The laser repetition period was 25 ns, divided into 256-time channels. Emission light, after passing through a bandpass filter (535/50 nm for $JF_{525}$; 585/40 nm for TMR; and 731/130 nm for $JF_{635}$, $JF_{646}$, and $JFX_{650}$, AVR Optics), was collected by an avalanche photodiode (SPCM-AQR-15, PerkinElmer) and analyzed by a digital frequency-domain module (DFD, FastFLIM, ISS). A 200 μm pinhole was used. Each image frame comprised 256 × 256 pixels (100 × 100 μm FOV) with a pixel dwell time of 50 μs. Photons from 6-8 frames were pooled for intensity and lifetime calculation, giving a sampling rate of 0.04-0.05 Hz.

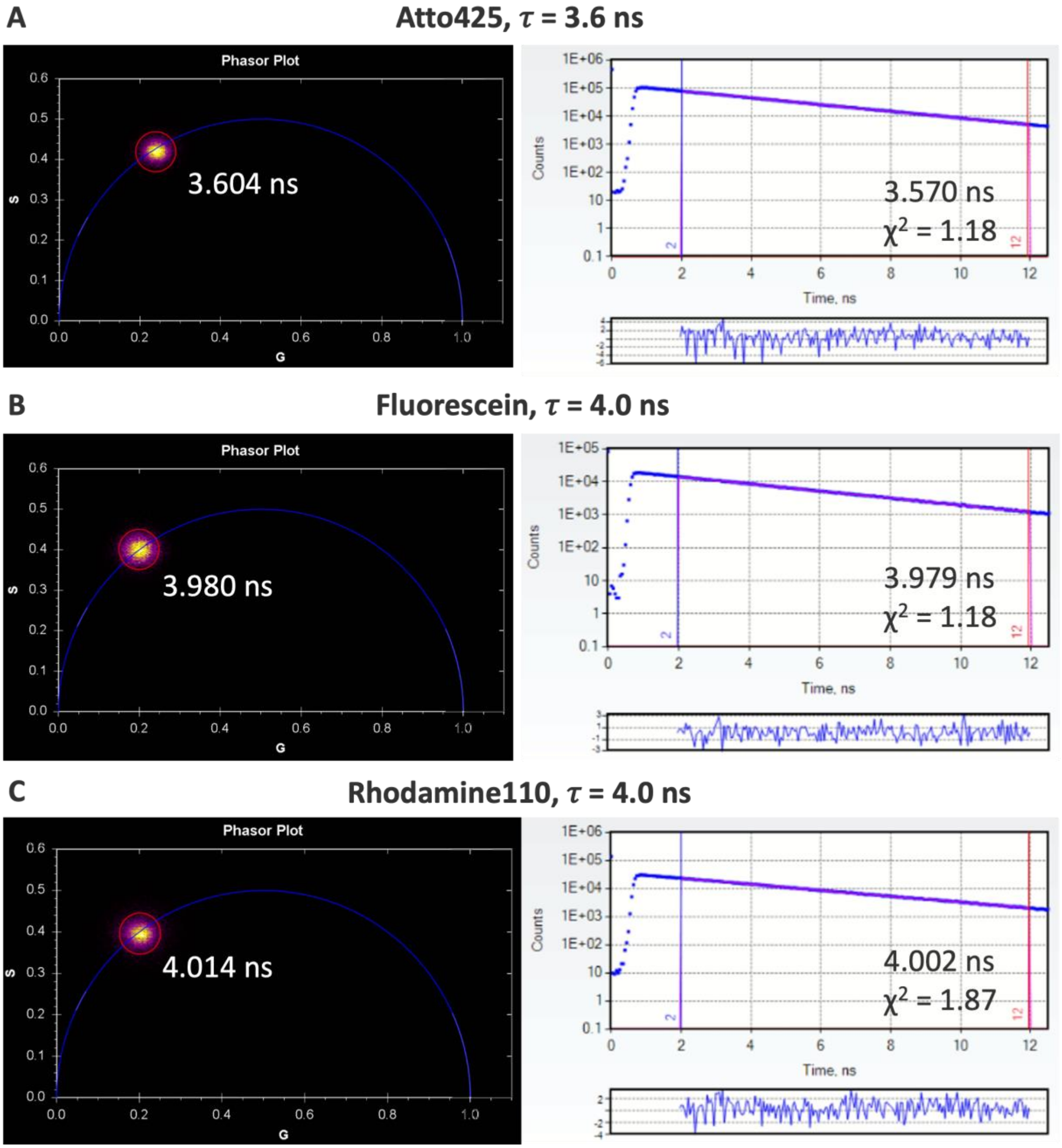


**Supplementary Figure S13. Comparison of fluorescence lifetimes obtained by digital frequency domain FastFLIM and time-correlated single-photon counting (TCSPC).** Representative phasor plots at 80 MHz (left) and fluorescence decay fits with residuals (right) for **(A)** Atto425 with known lifetime of 3.6 ns, **(B)** fluorescein, and **(C)** rhodamine 110 with known lifetime of 4.0 ns. Lifetimes obtained by DFD-FLIM are comparable with measurements from the TCSPC. Samples were excited by 2-photon laser (Insight X3, Spectra Physics) at 800 nm with a repetition rate of 80 MHz.

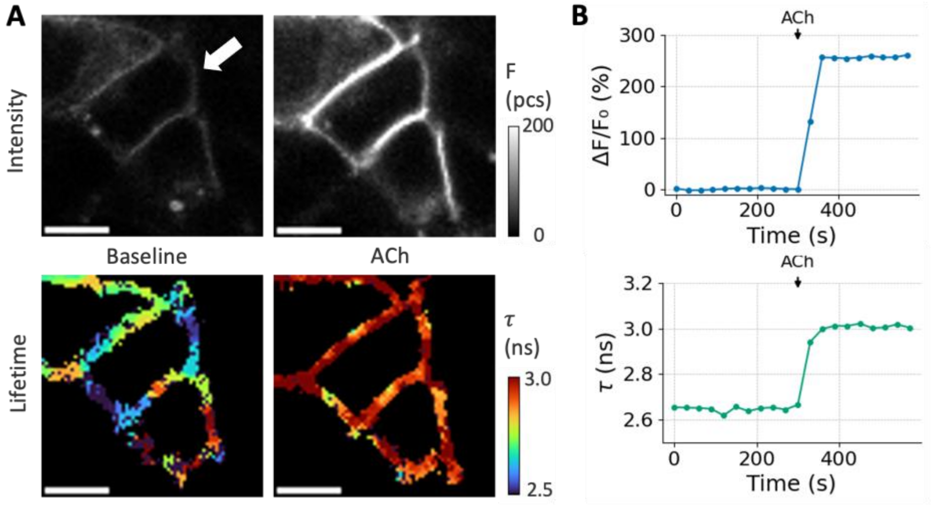


**Supplementary Figure S14. Our system was validated with a published acetylcholine sensor that exhibited lifetime increases upon ligand binding $\mathbf{GRAB_{ACh3.0}}$ in HEK293T cells.** Representative intensity (top row) and fluorescence lifetime (bottom row) images of HEK293T cells expressing $\mathbf{GRAB_{ACh3.0}}$. Cells were imaged at baseline, after addition of 100 µM acetylcholine (ACh). Scale bars: 10 µm. Intensity was reported as photon counts (pc). **(B)** Representative time traces showing the reversibility of the sensor response in both fluorescence intensity ($\Delta F/F_0$, top) and lifetime ($\tau$, bottom) upon sequential addition of ACh. Time traces represent the response of highlighted cells in **(A)**. Responses agreed with previously published data[7–9].

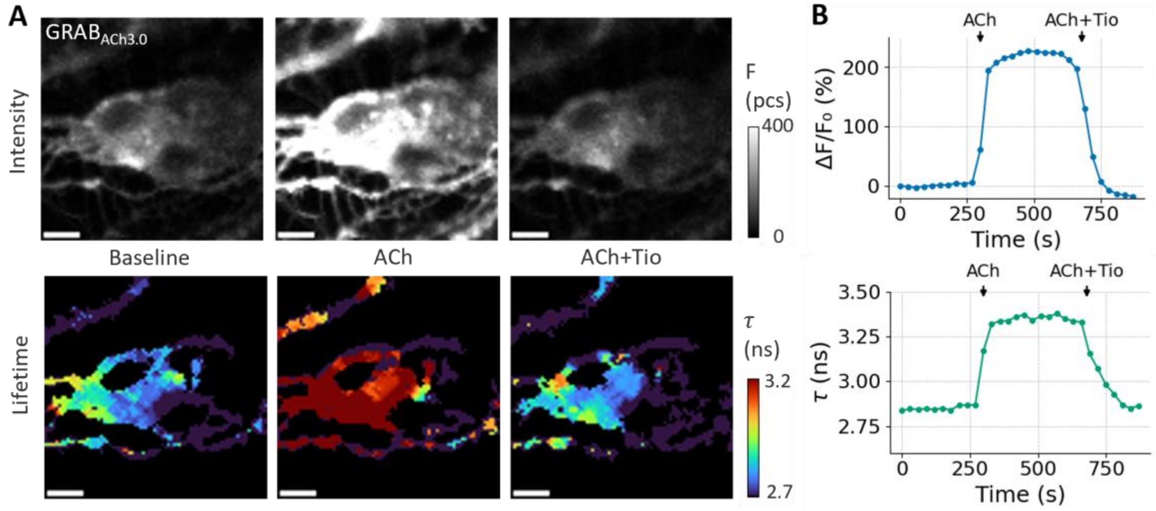


**Supplementary Figure S15. Our system was validated with a published acetylcholine sensor that exhibited lifetime increases upon ligand binding GRAB$_{ACh3.0}$ in primary neurons.** Representative intensity (top row) and fluorescence lifetime (bottom row) images of primary neurons expressing **GRAB$_{ACh3.0}$**. Cells were imaged at baseline, after addition of 100 µM acetylcholine (ACh). Scale bars: 10 µm. Intensity was reported as photon counts (pc). **(B)** Representative time traces showing the reversibility of the sensor response in both fluorescence intensity ($\Delta F/F_0$, top) and lifetime ($\tau$, bottom) upon sequential addition of ACh. The change was reversed by the co-application of the muscarinic ACh receptor (mAChR) antagonist tiotropium (Tio, 5 µM). Time traces represent the response of highlighted cells in **(A)**. Responses agreed with previously published data[7–9].

**Supplementary Table 1. Photophysical properties of rhodamine-based dyes.** $\lambda_{abs}$, absorption maximum; $\lambda_{em}$, emission maximum; ε, extinction coefficient; ϕ, quantum yield; $K_{L-Z}$, Lactone (L) – Zwitterion (Z) equilibrium

| Dye | $\lambda_{abs}$ (nm) | $\lambda_{em}$ (nm) | ε ($M^{-1}.cm^{-1}$) | Φ | $K_{L-Z}$[c] | Reference |
|---|---|---|---|---|---|---|
| $JF_{525}$ | 525 | 549 | 122,000[a] | 0.91 | 0.068 | [10] |
| TMR | 548 | 572 | 78,000[b] | 0.41 | - | [11] |
| $JF_{635}$ | 635 | 652 | 167,000[a] | 0.56 | <0.0001 | [10] |
| $JF_{646}$ | 646 | 664 | 152,000[a] | 0.54 | 0.0012 | [10] |
| JFX650 | 650 | 667 | >150,000[a] | 0.53 | 0.015 | [12] |

[a]Measured in ethanol or TFE with 0.1% TFA.

[b]Measured in 10 mM HEPES, pH 7.3.

[c]Equilibrium constant measured in 1:1 (vol:vol) dioxane:water.

**Supplementary Table 2. Mutations and linkers used for all variants in this study.** Sequence comparison to dHaloLife where - indicating no difference to dHaloLife. A total of 42 sensors were tested in this study.

| **Variant** | **Dye** | **DRD1 – N** | **N linker (NL)** | **cpHaloTag** | **C linker (CL)** | **DRD1 – C** | **Results** |
|---|---|---|---|---|---|---|---|
| Variant 0 | $JF_{635}$ | - | LSSLI | - | NHDQL | - | Poor membrane localization |
| Variant 1 | $JF_{525}$<br>TMR<br>$JF_{635}$<br>$JF_{646}$<br>$JFX_{650}$ | - | PK | - | PFAR | - | Summarized in **Supplementary Table 4** |
| Variant 2 | $JF_{525}$<br>TMR<br>$JF_{635}$<br>$JF_{646}$<br>$JFX_{650}$ | - | GG | - | PEFAR | - | Summarized in **Supplementary Table 4** |
| Variant 2.1 | $JF_{646}$<br>$JFX_{650}$ | - | LSSLI | - | PEFARNHDQL | - | Poor membrane localization |
| Variant 3 (dHaloLife) | $JF_{525}$<br>TMR<br>$JF_{635}$<br>$JF_{646}$<br>$JFX_{650}$ | - | (none) | - | PEFAR | - | Summarized in **Supplementary Table 4** |
| dHaloLife-mut | $JF_{525}$<br>TMR<br>$JF_{635}$<br>$JF_{646}$<br>$JFX_{650}$ | D103A | (none) | - | PEFAR | - | Summarized in **Supplementary Table 4** |
| Variant 3.1 | $JF_{635}$ | - | PEFAR | - | (none) | - | Poor membrane localization |
| Variant 3.2 | $JF_{635}$ | - | (none) | - | (none) | - | Poor membrane localization |
| Variant 4 | $JF_{525}$<br>TMR<br>$JF_{635}$<br>$JF_{646}$ | - | (none) | cpHT9 | PEFAR | - | Summarized in **Supplementary Table 4** |

| | | | | | | | |
|---|---|---|---|---|---|---|---|
| | $JFX_{650}$ | | | | | | |
| Variant 4.1 | $JF_{635}$ $JF_{646}$ $JFX_{650}$ | - | (none) | cpHT10 | PEFAR | - | Brightness is too low |
| Variant 5.0 | $JF_{525}$ TMR $JF_{635}$ $JF_{646}$ | F129A Insert 224Q | (none) | - | - | - | Poor membrane localization |
| Variant 5 | $JF_{525}$ TMR $JF_{635}$ $JF_{646}$ $JFX_{650}$ | I217V | (none) | - | PEFAR | V544L | Summarized in **Supplementary Table 4** |

**Supplementary Table 3. Mean and s.e.m. of fluorescence intensity and lifetime of each variant-dye combination.**

| Variant | Dye | Sensor number | Intensity change ($\Delta F/F_0$, %) | Lifetime change ($\Delta\tau$, ns) | Baseline lifetime ($\tau$, ns) | Sample size |
|---|---|---|---|---|---|---|
| Variant 1 | $JF_{525}$ | Sensor #1 | -11.8 ± 0.7 | -0.129 ± 0.005 | 4.126 ± 0.1183 | 15 cells 3 independent experiments |
| | TMR | Sensor #2 | -0.6 ± 1.1 | 0.126 ± 0.003 | 3.642 ± 0.153 | 15 cells 3 independent experiments |
| | $JF_{635}$ | Sensor #3 | -3.7 ± 1.7 | -0.005 ± 0.002 | 4.150 ± 0.049 | 15 cells 3 independent experiments |
| | $JF_{646}$ | Sensor #4 | 5.2 ± 1.7 | 0.057 ± 0.003 | 3.877 ± 0.029 | 15 cells 3 independent experiments |
| | $JFX_{650}$ | Sensor #5 | 10.7 ± 3.0 | 0.065 ± 0.007 | 4.370 ± 0.036 | 15 cells 3 independent experiments |
| Variant 2 | $JF_{525}$ | Sensor #6 | -7.3 ± 2.3 | -0.125 ± 0.013 | 4.027 ± 0.058 | 15 cells 3 independent experiments |
| | TMR | Sensor #7 | -4.6 ± 2.7 | -0.084 ± 0.021 | 3.797 ± 0.031 | 13 cells 3 independent experiments |
| | $JF_{635}$ | Sensor #8 | 3.2 ± 1.6 | -0.037 ± 0.006 | 4.305 ± 0.022 | 15 cells 3 independent experiments |
| | $JF_{646}$ | Sensor #9 | 4.1 ± 2.1 | -0.006 ± 0.004 | 4.156 ± 0.065 | 15 cells 3 independent experiments |
| | $JFX_{650}$ | Sensor #10 | 2.0 ± 3.7 | 0.000 ± 0.005 | 4.628 ± 0.041 | 15 cells 3 independent experiments |
| **Variant 3 (dHaloLife**) | $JF_{525}$ | Sensor #11 | 3.1 ± 0.9 | -0.026 ± 0.011 | 3.854 ± 0.039 | 15 cells 3 independent experiments |
| | TMR | Sensor #12 | -2.5 ± 1.6 | -0.179 ± 0.021 | 3.575 ± 0.069 | 19 cells 4 independent experiments |

| | | | | | | |
|---|---|---|---|---|---|---|
| | **$JF_{635}$** | Sensor #13 | **-34.3 ± 1.2** | -0.212 ± 0.004 | 4.586 ± 0.013 | **81 cells 16 independent experiments** |
| | $JF_{646}$ | Sensor #14 | 9.2 ± 7.8 | -0.080 ± 0.004 | 4.331 ± 0.047 | 12 cells 3 independent experiments |
| | $JFX_{650}$ | Sensor #15 | 12.5 ± 3.8 | -0.023 ± 0.006 | 4.556 ± 0.013 | 15 cells 3 independent experiments |
| Variant 4 | $JF_{525}$ | Sensor #16 | 4.1 ± 2.0 | 0.010 ± 0.008 | 3.397 ± 0.033 | 15 cells 3 independent experiments |
| | TMR | Sensor #17 | -6.0 ± 1.3 | -0.072 ± 0.012 | 3.358 ± 0.017 | 15 cells 3 independent experiments |
| | $JF_{635}$ | Sensor #18 | -7.9 ± 6.0 | -0.058 ± 0.009 | 4.320 ± 0.022 | 15 cells 3 independent experiments |
| | $JF_{646}$ | Sensor #19 | 2.2 ± 5.2 | -0.024 ± 0.008 | 4.204 ± 0.059 | 15 cells 3 independent experiments |
| | $JFX_{650}$ | Sensor #20 | 8.6 ± 2.9 | -0.066 ± 0.007 | 4.199 ± 0.038 | 15 cells 3 independent experiments |
| Variant 5 | $JF_{525}$ | Sensor #21 | 3.1 ± 2.1 | -0.086 ± 0.010 | 3.459 ± 0.070 | 15 cells 3 independent experiments |
| | TMR | Sensor #22 | -5.6 ± 1.0 | -0.141 ± 0.010 | 3.351 ± 0.071 | 15 cells 3 independent experiments |
| | $JF_{635}$ | Sensor #23 | -10.5 ± 3.3 | -0.117 ± 0.008 | 4.572 ± 0.087 | 20 cells 4 independent experiments |
| | $JF_{646}$ | Sensor #24 | 6.6 ± 3.9 | -0.048 ± 0.004 | 3.999 ± 0.039 | 15 cells 3 independent experiments |
| | $JFX_{650}$ | Sensor #25 | 12.5 ± 2.0 | -0.047 ± 0.006 | 4.296 ± 0.025 | 15 cells 3 independent experiments |
| dHaloLife-mut | $JF_{525}$ | - | -0.9 ± 0.8 | 0.033 ± 0.012 | 3.708 ± 0.068 | 15 cells 3 independent experiments |
| | TMR | - | -1.0 ± 0.5 | 0.014 ± 0.005 | 3.581 ± 0.107 | 15 cells |

| | | | | | |
|---|---|---|---|---|---|
| | | | | | 3 independent experiments |
| $JF_{635}$ | - | -1.0 ± 1.0 | 0.013 ± 0.004 | 4.576 ± 0.084 | 20 cells 4 independent experiments |
| $JF_{646}$ | - | -3.3 ± 1.6 | -0.013 ± 0.002 | 4.281 ± 0.038 | 15 cells 3 independent experiments |
| $JFX_{650}$ | - | -0.8 ± 1.7 | 0.035 ± 0.007 | 4.682 ± 0.016 | 20 cells 4 independent experiments |

**Supplementary Table 4. 95% confidence interval of mean of each variant-dye combination.**

| Variant | Dye | Sensor number | Lower 95% CI of mean intensity change (%) | Upper 95% CI of mean intensity change (%) | Lower 95% CI of mean lifetime change (ns) | Upper 95% CI of mean lifetime change (ns) | Sample size |
|---|---|---|---|---|---|---|---|
| Variant 1 | $JF_{525}$ | Sensor #1 | -13.3 | -10.3 | -0.141 | -0.118 | 15 cells 3 independent experiments |
| | TMR | Sensor #2 | -2.9 | 1.7 | 0.119 | 0.133 | 15 cells 3 independent experiments |
| | $JF_{635}$ | Sensor #3 | -7.4 | -0.01 | -0.010 | 0.000 | 15 cells 3 independent experiments |
| | $JF_{646}$ | Sensor #4 | 1.5 | 8.9 | 0.052 | 0.063 | 15 cells 3 independent experiments |
| | $JFX_{650}$ | Sensor #5 | 4.4 | 17.1 | 0.050 | 0.079 | 15 cells 3 independent experiments |
| Variant 2 | $JF_{525}$ | Sensor #6 | -12.2 | -2.3 | -0.153 | -0.098 | 15 cells 3 independent experiments |
| | TMR | Sensor #7 | -10.3 | 1.2 | -0.128 | -0.039 | 13 cells 3 independent experiments |
| | $JF_{635}$ | Sensor #8 | -0.3 | 6.7 | -0.050 | -0.024 | 15 cells 3 independent experiments |
| | $JF_{646}$ | Sensor #9 | -0.5 | 8.7 | -0.015 | 0.002 | 15 cells 3 independent experiments |

| | | | | | | | |
|---|---|---|---|---|---|---|---|
| | $JFX_{650}$ | Sensor #10 | -6.0 | 9.9 | -0.010 | 0.010 | 15 cells 3 independent experiments |
| **Variant 3 (dHaloLife)** | $JF_{525}$ | Sensor #11 | 1.2 | 5.0 | -0.050 | -0.003 | 15 cells 3 independent experiments |
| | TMR | Sensor #12 | -5.8 | 0.9 | -0.222 | -0.136 | 19 cells 4 independent experiments |
| | **$JF_{635}$** | **Sensor #13** | **-36.6** | **-31.9** | **-0.219** | **-0.205** | **81 cells 16 independent experiments** |
| | $JF_{646}$ | Sensor #14 | -7.9 | 26.3 | -0.089 | -0.072 | 12 cells 3 independent experiments |
| | $JFX_{650}$ | Sensor #15 | 4.4 | 20.7 | -0.035 | -0.012 | 15 cells 3 independent experiments |
| Variant 4 | $JF_{525}$ | Sensor #16 | -0.3 | 8.5 | -0.008 | 0.028 | 15 cells 3 independent experiments |
| | TMR | Sensor #17 | -8.8 | -3.2 | -0.098 | -0.047 | 15 cells 3 independent experiments |
| | $JF_{635}$ | Sensor #18 | -20.7 | 4.9 | -0.078 | -0.038 | 15 cells 3 independent experiments |
| | $JF_{646}$ | Sensor #19 | -9.0 | 13.5 | -0.040 | -0.007 | 15 cells 3 independent experiments |
| | $JFX_{650}$ | Sensor #20 | 2.4 | 14.8 | -0.082 | -0.05 | 15 cells 3 independent experiments |

| | | | | | | | |
|---|---|---|---|---|---|---|---|
| Variant 5 | $JF_{525}$ | Sensor #21 | -1.4 | 7.7 | -0.109 | -0.064 | 15 cells 3 independent experiments |
| | TMR | Sensor #22 | -7.9 | -3.4 | -0.163 | -0.120 | 15 cells 3 independent experiments |
| | $JF_{635}$ | Sensor #23 | -17.3 | -3.7 | -0.133 | -0.100 | 20 cells 4 independent experiments |
| | $JF_{646}$ | Sensor #24 | -1.7 | 14.9 | -0.056 | -0.040 | 15 cells 3 independent experiments |
| | $JFX_{650}$ | Sensor #25 | 8.2 | 16.8 | -0.059 | -0.035 | 15 cells 3 independent experiments |
| dHaloLife-mut | $JF_{525}$ | - | -2.6 | 0.7 | 0.008 | 0.057 | 15 cells 3 independent experiments |
| | TMR | - | -2.0 | 0.1 | 0.004 | 0.025 | 15 cells 3 independent experiments |
| | $JF_{635}$ | - | -3.1 | 1.1 | 0.004 | 0.022 | 20 cells 4 independent experiments |
| | $JF_{646}$ | - | -6.7 | 0.0 | -0.017 | -0.009 | 15 cells 3 independent experiments |
| | $JFX_{650}$ | - | -4.5 | 2.8 | 0.020 | 0.049 | 20 cells 4 independent experiments |

**Supplementary Table 5. Statistical analysis of the fluorescence intensity of dHaloLife$_{635}$ to co-application of antagonist SCH-23390 (Supplementary Figure S8).** One-way repeated-measures ANOVA with Tukey's multiple comparisons test. F (DFn, DFd) is the ANOVA statistic. Mean difference, 95% confidence interval of difference and adjusted p-values are reported for each comparison. n = 19 cells across 3 independent experiments.

| **One-way repeated-measures ANOVA** | |
|---|---|
| F (DFn, DFd) | F (2, 36) = 15.79 |
| P value | <0.0001 |
| P value summary | **** |
| Is there significant matching (P < 0.05)? | Yes |

| **Tukey's multiple comparisons test** | | |
|---|---|---|
| Mean diff. | Baseline vs. DA | 17 |
| | Baseline vs. DA+SCH | 6 |
| | DA vs. DA+SCH | -11 |
| 95.00% CI of diff. | Baseline vs. DA | 9 to 24 |
| | Baseline vs. DA+SCH | -1 to 13 |
| | DA vs. DA+SCH | -18 to -3 |
| Summary with adjusted p-value | Baseline vs. DA | ****p<0.0001 |
| | Baseline vs. DA+SCH | ns p=0.1257 |
| | DA vs. DA+SCH | **p=0.0031 |

**Supplementary Table 6. Statistical analysis of the fluorescence lifetime of dHaloLife$_{635}$ to co-application of antagonist SCH-23390 (Supplementary Figure S8).** One-way repeated-measures ANOVA with Tukey's multiple comparisons test. F (DFn, DFd) is the ANOVA statistic. Mean difference, 95% confidence interval of difference and adjusted p-values are reported for each comparison. n = 19 cells across 3 independent experiments.

| **One-way repeated-measures ANOVA** | |
|---|---|
| F (DFn, DFd) | F (2, 36) = 287.7 |
| P value | <0.0001 |
| P value summary | **** |
| Is there significant matching (P < 0.05)? | Yes |

| **Tukey's multiple comparisons test** | | |
|---|---|---|
| Mean diff. | Baseline vs. DA | 0.199 |
| | Baseline vs. DA+SCH | -0.007 |
| | DA vs. DA+SCH | -0.206 |
| 95.00% CI of diff. | Baseline vs. DA | 0.175 to 0.223 |
| | Baseline vs. DA+SCH | -0.031 to 0.017 |
| | DA vs. DA+SCH | -0.230 to -0.182 |
| Summary with adjusted p-value | Baseline vs. DA | ****p<0.0001 |
| | Baseline vs. DA+SCH | ns p=0.7413 |
| | DA vs. DA+SCH | ****p<0.0001 |

**Supplementary Table 7. Statistical analysis of the fluorescence intensity response of dHaloLife$_{635}$ to structurally related neurochemicals in HEK293T cells (Figure 3D).** Brown-Forsythe and Welch ANOVA tests for the intensity ($\Delta F/F_0$) responses to 1 µM L-DOPA, GABA, NE, 5-HT and DA, with Dunnett's T3 multiple comparisons against DA. W (DFn, DFd) is the Welch ANOVA statistic. F (DFn, DFd) is the Brown-Forsythe statistic. Mean difference, 95% confidence interval of difference and adjusted p-values are reported for each comparison. n ≥ 14 cells across ≥ 3 independent experiments.

| **Brown-Forsythe ANOVA test** | |
|---|---|
| F* (DFn, DFd) | 22.24 (4.000, 66.64) |
| P value | <0.0001 |
| P value summary | **** |
| Significant diff. among means (P < 0.05)? | Yes |

| **Welch's ANOVA test** | |
|---|---|
| W (DFn, DFd) | 17.54 (4.000, 36.20) |
| P value | <0.0001 |
| P value summary | **** |
| Significant diff. among means (P < 0.05)? | Yes |

| **Dunnett's T3 multiple comparisons test** | | |
|---|---|---|
| Mean diff. | DA vs L-DOPA | -24.5 |
| | DA vs NE | -17.6 |
| | DA vs 5-HT | -24.0 |
| | DA vs GABA | -25.0 |
| 95.00% CI of diff. | DA vs L-DOPA | -33.9 to -15.2 |
| | DA vs NE | -28.8 to -6.3 |
| | DA vs 5-HT | -32.5 to -15.6 |
| | DA vs GABA | -33.80 to -16.1 |
| Summary with adjusted p-value | DA vs L-DOPA | ****p<0.0001 |
| | DA vs NE | **p=0.0012 |
| | DA vs 5-HT | ****p<0.0001 |
| | DA vs GABA | ****p<0.0001 |

**Supplementary Table 8. Statistical analysis of the fluorescence lifetime response of dHaloLife$_{635}$ to structurally related neurochemicals in HEK293T cells (Figure 3D).** Brown-Forsythe and Welch ANOVA tests for the lifetime (Δτ) responses to 1 µM L-DOPA, GABA, NE, 5-HT and DA, with Dunnett's T3 multiple comparisons against DA. W (DFn, DFd) is the Welch ANOVA statistic. F (DFn, DFd) is the Brown-Forsythe statistic. Mean difference, 95% confidence interval of difference and adjusted p-values are reported for each comparison. n ≥ 14 cells across ≥ 3 independent experiments.

| **Brown-Forsythe ANOVA test** | |
|---|---|
| F* (DFn, DFd) | 109.1 (4.000, 71.68) |
| P value | <0.0001 |
| P value summary | **** |
| Significant diff. among means (P < 0.05)? | Yes |

| **Welch's ANOVA test** | |
|---|---|
| W (DFn, DFd) | 76.94 (4.000, 36.39) |
| P value | <0.0001 |
| P value summary | **** |
| Significant diff. among means (P < 0.05)? | Yes |

| **Dunnett's T3 multiple comparisons test** | | |
|---|---|---|
| Mean diff. | DA vs L-DOPA | -0.080 |
| | DA vs NE | -0.061 |
| | DA vs 5-HT | -0.108 |
| | DA vs GABA | -0.106 |
| 95.00% CI of diff. | DA vs L-DOPA | -0.098 to -0.062 |
| | DA vs NE | -0.081 to -0.041 |
| | DA vs 5-HT | -0.128 to -0.089 |
| | DA vs GABA | -0.122 to -0.089 |
| Summary with adjusted p-value | DA vs L-DOPA | ****p<0.0001 |
| | DA vs NE | ****p<0.0001 |
| | DA vs 5-HT | ****p<0.0001 |
| | DA vs GABA | ****p<0.0001 |

**Supplementary Table 9. GPCR-based sensors for neurotransmitters and neuromodulators that exhibit lifetime change.**

| Sensor | Target | Lifetime change ($\Delta\tau$) | Protein scaffold | Emission color | Reference |
|---|---|---|---|---|---|
| $GRAB_{ACh3.0}$ | Acetylcholine | 0.17 – 0.37 ns | cpGFP | Green | [7], this paper |
| dLight3.8 | Dopamine | 0.240 ns | cpGFP | Green | [13] |
| dLight3.6 | Dopamine | 0.105 ns | cpGFP | Green | [13] |
| $dHaloLife_{635}$ | Dopamine | -0.21 ns | cpHaloTag | Far-red | This paper |

**Supplementary Table 10. Fluorescence intensity change, lifetime change, baseline lifetime, 95% confidence interval (CI) of intensity and lifetime changes of the $GRAB_{ACh3.0}$ using the digital frequency domain FLIM in Yeh Lab.** Data reported as mean ± s.e.m.

| **Sensor** | **Intensity change ($\Delta F/F_0$, %)** | **Lifetime change ($\Delta\tau$, ns)** | **Baseline lifetime ($\tau$, ns)** | **CI of intensity change (%)** | **CI of lifetime change (ns)** | **Sample size** |
|---|---|---|---|---|---|---|
| $GRAB_{ACh3.0}$ | 214.7 ± 19.3 | 0.365 ± 0.005 | 2.670 ± 0.007 | 173.2, 256.2 | 0.353, 0.376 | n = 15 cells from 3 independent experiments |

**Supplementary Table 11. Brightness comparison of dHaloLife to common fluorescent proteins.** ε, extinction coefficient; ϕ, quantum yield.

| Fluorophore | Φ | ε ($M^{-1}.cm^{-1}$) | Brightness (ε * Φ) *$10^3$ | Brightness comparison to HT7 – $JF_{635}$ | Reference |
|---|---|---|---|---|---|
| HaloTag7 – $JF_{635}$ | 0.675 | 167,000 | 112.73 | 1 | [14] |
| HaloTag9 – $JF_{635}$ | 0.67 | 167,000 | 111.89 | 1/0.99 | [14] |
| EGFP | 0.60 | 55,900 | 33.54 | 1/3.36 | [15,16] |
| mCherry | 0.22 | 72,000 | 15.84 | 1/7.12 | [15,16] |

**Supplementary Table 12. dHaloLife DNA sequence.** Nucleotide sequence of the dHaloLife coding region. The total length is 2163 nucleotides. Color-coding designates the N-terminal domain of the hDRD1 receptor sequence (gray), followed by the C-terminal domain of the HaloTag (yellow), a flexible linker connecting the 2 domains of the HaloTag, then the N-terminal domain of the HaloTag (yellow), creating a cpHaloTag, then C-linker of the sensor (black) followed by the C-terminal domain of the hDRD1 (gray).

| | | |
|---|---|---|
| 1 | ATGAGGACTCTGAACACCTCTGCCATGGAC | 30 |
| 31 | GGGACTGGGCTGGTGGTGGAGAGGGACTTC | 60 |
| 61 | TCTGTTCGTATCCTCACTGCCTGTTTCCTG | 90 |
| 91 | TCGCTGCTCATCCTGTCCACGCTCCTGGGG | 120 |
| 121 | AACACGCTGGTCTGTGCTGCCGTTATCAGG | 150 |
| 151 | TTCCGACACCTGCGGTCCAAGGTGACCAAC | 180 |
| 181 | TTCTTTGTCATCTCCTTGGCTGTGTCAGAT | 210 |
| 211 | CTCTTGGTGGCCGTCCTGGTCATGCCCTGG | 240 |
| 241 | AAGGCAGTGGCTGAGATTGCTGGCTTCTGG | 270 |
| 271 | CCCTTTGGGTCCTTCTGTAACATCTGGGTG | 300 |
| 301 | GCCTTTGACATCATGTGCTCCACTGCATCC | 330 |
| 331 | ATCCTCAACCTCTGTGTGATCAGCGTGGAC | 360 |
| 361 | AGGTATTGGGCTATCTCCAGCCCTTTCCGG | 390 |
| 391 | TATGAGAGAAAGATGACCCCCAAGGCAGCC | 420 |
| 421 | TTCATCCTGATCAGTGTGGCATGGACCTTG | 450 |
| 451 | TCTGTACTCATCTCCTTCATCCCAGTGCAG | 480 |
| 481 | CTCAGCTGGCACAAGGCAAAACCCACAAGC | 510 |
| 511 | CCCTCTGATGGAAATGCCACTTCCCTGGCT | 540 |
| 541 | GAGACCATAGACAACTGTGACTCCAGCCTC | 570 |
| 571 | AGCAGGACATATGCCATCTCATCCTCTGTA | 600 |
| 601 | ATCAGCTTTTACATCCCTGTGGCCATCATG | 630 |
| 631 | ATTGTCACCTACACCAGGATCTACAGGATT | 660 |
| 661 | GCTCAGAAAGAGACCTTCCAGGCCTTCCGC | 690 |
| 691 | ACCACCGACGTCGGCCGCAAGCTGATCATC | 720 |
| 721 | GATCAGAACGTTTTTATCGAGGGTACGCTG | 750 |
| 751 | CCGATGGGTGTCGTCCGCCCGCTGACTGAA | 780 |
| 781 | GTCGAGATGGACCATTACCGCGAGCCGTTC | 810 |
| 811 | CTGAATCCTGTTGACCGCGAGCCACTGTGG | 840 |
| 841 | CGCTTCCCAAACGAGCTGCCAATCGCCGGT | 870 |
| 871 | GAGCCAGCGAACATCGTCGCGCTGGTCGAA | 900 |
| 901 | GAATACATGGACTGGCTGCACCAGTCCCCT | 930 |
| 931 | GTCCCGAAGCTGCTGTTCTGGGGCACCCCA | 960 |
| 961 | GGCGTTCTGATCCCACCGGCCGAAGCCGCT | 990 |
| 991 | CGCCTGGCCAAAAGCCTGCCTAACTGCAAG | 1020 |

| | | |
|---|---|---|
| 1021 | GCTGTGGACATCGGCCCGGGTCTGAATCTG | 1050 |
| 1051 | CTGCAAGAAGACAACCCGGACCTGATCGGC | 1080 |
| 1081 | AGCGAGATCGCGCGCTGGCTGTCGACGCTC | 1110 |
| 1111 | GAGATTTCCGGCGGAGGAACAGGTGGTTCT | 1140 |
| 1141 | GGTGGAACAGGGGGTAGCGGAGGTACAGGA | 1170 |
| 1171 | GGAAGTATGGCGGAGATCGGAACTGGATTC | 1200 |
| 1201 | CCGTTTGATCCGCATTATGTGGAAGTTCTG | 1230 |
| 1231 | GGAGAGCGCATGCATTATGTGGACGTTGGT | 1260 |
| 1261 | CCTCGTGATGGGACACCAGTGCTGTTCCTT | 1290 |
| 1291 | CACGGCAATCCGACATCGTCGTACGTGTGG | 1320 |
| 1321 | CGTAATATCATCCCGCACGTTGCCCCCACG | 1350 |
| 1351 | CACCGCTGCATTGCCCCTGACTTAATTGGT | 1380 |
| 1381 | ATGGGGAAAAGTGATAAGCCTGATCTGGGG | 1410 |
| 1411 | TACTTCTTTGACGACCACGTACGCTTCATG | 1440 |
| 1441 | GATGCTTTTATTGAAGCATTGGGTTTGGAG | 1470 |
| 1471 | GAAGTAGTTTTGGTGATCCATGATTGGGGT | 1500 |
| 1501 | AGTGCTCTGGGGTTCCATTGGGCCAAGCGT | 1530 |
| 1531 | AACCCAGAACGCGTGAAAGGAATTGCCTTT | 1560 |
| 1561 | ATGGAGTTCATCCGTCCGATTCCAACATGG | 1590 |
| 1591 | GACGAATGGCCAGAATTTGCACGCAAAAGA | 1620 |
| 1621 | GAAACTAAAGTCCTGAAGACTCTGTCGGTG | 1650 |
| 1651 | ATCATGGGTGTGTTTGTGTGCTGTTGGCTA | 1680 |
| 1681 | CCTTTCTTCATCTTGAACTGCATTTTGCCC | 1710 |
| 1711 | TTCTGTGGGTCTGGGGAGACGCAGCCCTTC | 1740 |
| 1741 | TGCATTGATTCCAACACCTTTGACGTGTTT | 1770 |
| 1771 | GTGTGGTTTGGGTGGGCTAATTCATCCTTG | 1800 |
| 1801 | AACCCCATCATTTATGCCTTTAATGCTGAT | 1830 |
| 1831 | TTTCGGAAGGCATTTTCAACCCTCTTAGGA | 1860 |
| 1861 | TGCTACAGACTTTGCCCTGCGACGAATAAT | 1890 |
| 1891 | GCCATAGAGACGGTGAGTATCAATAACAAT | 1920 |
| 1921 | GGGGCCGCGATGTTTTCCAGCCATCATGAG | 1950 |
| 1951 | CCACGAGGCTCCATCTCCAAGGAGTGCAAT | 1980 |
| 1981 | CTGGTTTACCTGATCCCACATGCTGTGGGC | 2010 |
| 2011 | TCCTCTGAGGACCTGAAAAAGGAGGAGGCA | 2040 |
| 2041 | GCTGGCATCGCCAGACCCTTGGAGAAGCTG | 2070 |
| 2071 | TCCCCAGCCCTATCGGTCATATTGGACTAT | 2100 |
| 2101 | GACACTGACGTCTCTCTGGAGAAGATCCAA | 2130 |
| 2131 | CCCATCACACAAAACGGTCAGCACCCAACC | 2160 |
| 2161 | TGA | 2163 |

dHaloLife$_{635}$ in HEK293T cells

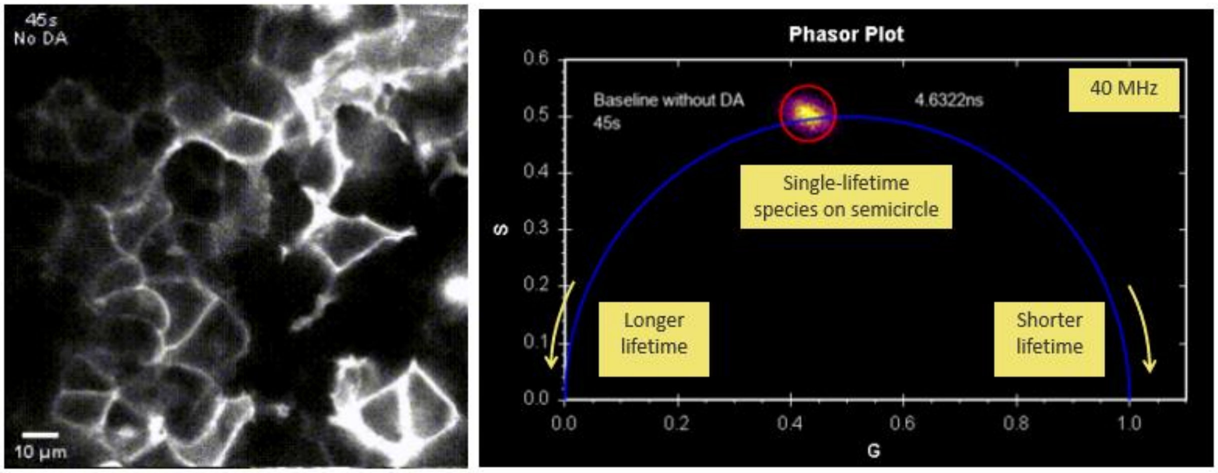


**Supplementary Video 1. Time-lapse fluorescence lifetime imaging and phasor analysis of dHaloLife$_{635}$ in HEK293T cells.** Dynamic imaging and phasor plot analysis of dHaloLife$_{635}$ in HEK293T cells. The video displays two synchronized panels: a time-lapse fluorescence intensity view showing HEK293T cells expressing dHaloLife$_{635}$ (scale bar = 10 µm) and a corresponding phasor plot tracking lifetime changes. The phasor plot is configured at 40 MHz, mapping single-lifetime species on the universal semicircle (ranging from shorter to longer lifetimes). The response to dopamine co-applied with DRD1-specific antagonist SCH-23390 (SCH) reverted the intensity and lifetime changes back to the baseline.

dHaloLife-mut$_{635}$ in HEK293T cells

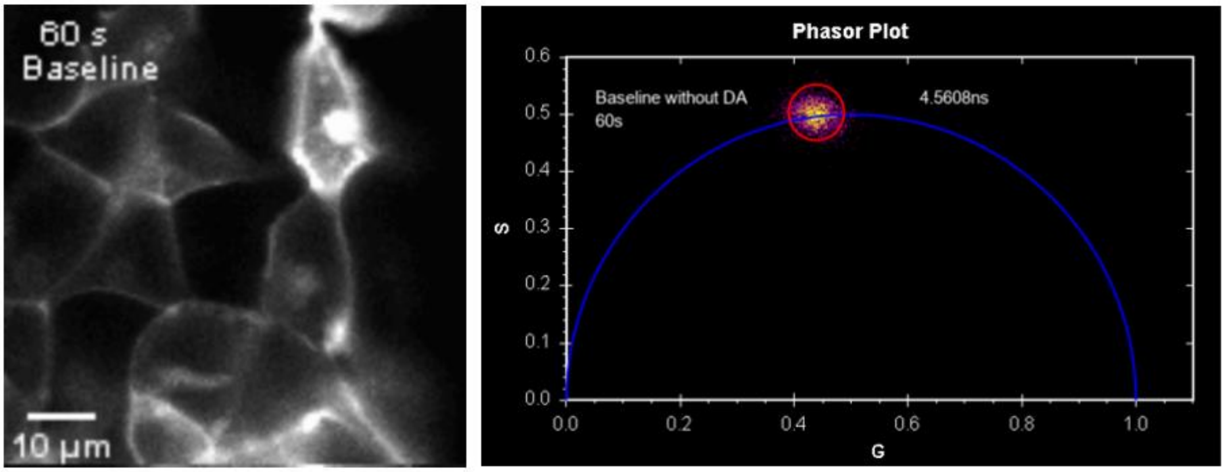


**Supplementary Video 2. Time-lapse fluorescence lifetime imaging and phasor analysis of dHaloLife-mut$_{635}$ in HEK293T cells.** Dynamic imaging and phasor plot analysis of dHaloLife-mut$_{635}$ in HEK293T cells. The video displays two synchronized panels: a time-lapse fluorescence intensity view showing HEK293T cells expressing dHaloLife-mut$_{635}$ (scale bar = 10 μm) and a corresponding phasor plot tracking insignificant lifetime changes with dopamine added. The phasor plot is configured at 40 MHz, mapping single-lifetime species on the universal semicircle.

dHaloLife$_{635}$ in primary neurons

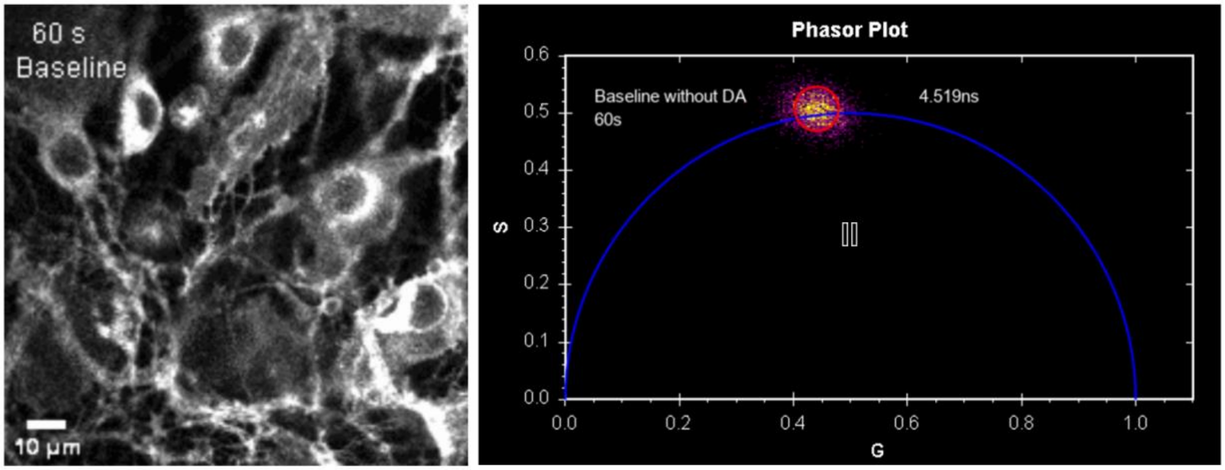


**Supplementary Video 3. Time-lapse fluorescence lifetime imaging and phasor analysis of dHaloLife$_{635}$ in primary neurons.** Dynamic imaging and phasor plot analysis of dHaloLife$_{635}$ in primary neurons. The video features side-by-side visualization of primary neurons expressing dHaloLife$_{635}$ (scale bar = 10 µm) alongside a real-time phasor plot. The phasor distribution shifts along the 40 MHz universal semicircle. The response to dopamine co-applied with DRD1-specific antagonist SCH-23390 (SCH) reverted the intensity and lifetime changes back to the baseline.

**References**


1. Digman, M. A., Caiolfa, V. R., Zamai, M. & Gratton, E. The Phasor Approach to Fluorescence Lifetime Imaging Analysis. *Biophysical Journal* **94**, L14–L16 (2008).
2. Redford, G. I. & Clegg, R. M. Polar Plot Representation for Frequency-Domain Analysis of Fluorescence Lifetimes. *J Fluoresc* **15**, 805–815 (2005).
3. Mirdita, M. *et al.* ColabFold: Making Protein folding accessible to all. *Nature Methods* https://doi.org/10.1038/s41592-022-01488-1 (2022) doi:10.1038/s41592-022-01488-1.
4. Jumper, J. *et al.* Highly accurate protein structure prediction with AlphaFold. *Nature* https://doi.org/10.1038/s41586-021-03819-2 (2021) doi:10.1038/s41586-021-03819-2.
5. Meng, E. C. *et al.* UCSF ChimeraX: Tools for structure building and analysis. *Protein Science* **32**, e4792 (2023).
6. Imoto, H. himoto/hillfit. (2026).
7. Ma, P. *et al.* Fast and slow: Recording neuromodulator dynamics across both transient and chronic time scales. *Science Advances* **10**, eadi0643 (2024).
8. Jing, M. *et al.* An optimized acetylcholine sensor for monitoring in vivo cholinergic activity. *Nat Methods* **17**, 1139–1146 (2020).
9. Nguyen, A.-T. *et al.* Fluorescence-lifetime-modulating probes for neural activity sensing. in *Reporters, Contrast Agents, and Molecular Probes for Biomedical Applications XVI* vol. 13339 6 (SPIE, 2025).
10. Grimm, J. B. *et al.* A general method to fine-tune fluorophores for live-cell and in vivo imaging. *Nat Methods* **14**, 987–994 (2017).

11. Grimm, J. B. *et al.* A general method to improve fluorophores for live-cell and single-molecule microscopy. *Nat Methods* **12**, 244–250 (2015).

12. Grimm, J. B. *et al.* A General Method to Improve Fluorophores Using Deuterated Auxochromes. *JACS Au* **1**, 690–696 (2021).

13. Tian, L. *et al.* Sensitive dLight for imaging broad-spectrum dopamine events across brain regions. Preprint at https://doi.org/10.21203/rs.3.rs-7313638/v1 (2025).

14. Frei, M. S. *et al.* Engineered HaloTag variants for fluorescence lifetime multiplexing. *Nat Methods* **19**, 65–70 (2022).

15. Lambert, T. J. FPbase: a community-editable fluorescent protein database. *Nat Methods* **16**, 277–278 (2019).

16. Zheng, Y. *et al.* In vivo multiplex imaging of dynamic neurochemical networks with designed far-red dopamine sensors. *Science* **388**, eadt7705 (2025).
